# Hippocampal subfield macrostructure and myelination demonstrate distinct maturational signatures

**DOI:** 10.64898/2026.09.07.749954

**Authors:** Kassandra Roger, Emily S. Nichols, Mahmoud Salman, Ali R. Khan, Emma G. Duerden

## Abstract

The hippocampus is essential for learning and memory and undergoes prolonged development throughout childhood and adolescence. Evidence concerning hippocampal maturation is based primarily on volumetric measures. However, macrostructural growth and myelination processes may represent distinct biological processes. Using Hippunfold, we quantified hippocampal subfield volumes and myelin-sensitive tissue concentrations as proxies for tissue maturation using T1w/T2w ratios in 175 community-recruited participants aged 9 months to 21 years. Hippocampal subfield volumes were best described by quadratic developmental trajectories, whereas T1w/T2w ratios increased linearly with age across all subfields. Males exhibited larger hippocampal volumes, and females demonstrated higher and steeper age-related increases in T1w/T2w ratios. Sex-related hippocampal subfield differences were observed despite largely comparable cognitive and behavioural profiles. Hippocampal subfield volumes and myelin follow distinct developmental pathways, revealing sex-specific patterns of development that are not fully captured by morphology alone.

## 1. Introduction

The hippocampus is a brain structure located in the medial temporal lobe comprising several subfields including the cornu ammonis 1 to 4 (CA1-4), the dentate gyrus (DG) and the subiculum (Cajal, 1901; Lorente de Nó, 1934). It has attracted considerable interest in the neuroscience literature, primarily because of its role in memory and learning, spatial navigation (Deshmukh & Knierim, 2012) and emotion (Nichols et al., 2023). Various developmental psychiatric and neurological conditions are also known to affect the hippocampus, such as schizophrenia (Nakahara et al., 2018; van Erp et al., 2016), depression (Schmaal et al., 2016; Sun et al., 2023), bipolar disorder (Haukvik et al., 2022; Hibar et al., 2016), autism spectrum disorder (ASD; Deniz et al., 2026), and attention-deficit/hyperactivity disorder (ADHD; Hoogman et al., 2017). Consequently, understanding hippocampal development and sex differences in developmental trajectories has become a major focus of developmental neuroscience, particularly given that many of these disorders emerge during childhood and adolescence and are sexually dimorphic. To date, research has largely focused on characterizing age-related differences in hippocampal morphology, most commonly through volumetric measures. However, developmental processes underlying hippocampal maturation, such as myelination, remain comparatively understudied despite their potential importance for cognitive development and vulnerability to early life stress (Nichols et al., 2025).

A substantial body of research has examined age-related changes in hippocampal subfield volumes in typical and atypical childhood and adolescent development. However, findings remain inconsistent, with studies reporting positive, negative, nonlinear, or absent associations between age and individual subfield volumes (Karat et al., 2024; Krogsrud et al., 2014; Lee et al., 2014; Mu et al., 2020; Sussman et al., 2016; Tamnes et al., 2018). A recent meta-analysis of 11 developmental studies concluded that age was positively associated with DG and CA3-4 volumes across childhood and adolescence, but not with subiculum or CA1 volumes (Homayouni et al., 2023).

Hippocampal volumes also appear to vary by sex, but results are still inconsistent, with early studies reporting larger volumes in females and more recent studies reporting larger volumes in males or no sex differences at all (Karat et al., 2024; Krogsrud et al., 2014; Mu et al., 2020; Riggins et al., 2018; Sussman et al., 2016; Tamnes et al., 2018). This is in the context of a well-established sexual dimorphism in brain development, with males, on average, having larger brains than females (Ruigrok et al., 2014). Thus, volumetric measures may not be optimally suited to capture the dominant neurodevelopmental processes occurring during this period. While the basic architecture of the hippocampus is established prenatally, tissue maturation continues throughout childhood and adolescence through processes such as myelination and circuit refinement (Ábrahám et al., 2010; Lavenex & Banta Lavenex, 2013; Seress & Ábrahám, 2008). White matter development follows a prolonged trajectory extending well into adulthood, whereas many aspects of grey matter development occur substantially earlier (Duerden et al., 2020; Houston et al., 2014; Mills et al., 2016). Consequently, measures of tissue composition may provide greater sensitivity to ongoing developmental change than morphology alone.

Recent advances in hippocampal segmentation have created new opportunities to characterize morphological and microstructural development (DeKraker et al., 2022). Hippunfold is a surface-based segmentation framework. In addition to improving anatomical precision (DeKraker et al., 2018), Hippunfold enables the extraction of biologically informative measures such as T1w/T2w ratios, proxies for tissue myelination (Ganzetti et al., 2014). This is particularly relevant during childhood and adolescence, when ongoing myelination represents a major component of brain maturation (Houston et al., 2014; Mills et al., 2016; Norbom et al., 2020). Consequently, combining volumetric and myelin-sensitive measures may provide a more comprehensive understanding of hippocampal development than either measure alone.

The primary aim of this study was to characterize age-related variations in hippocampal subfield morphology and tissue maturation derived from T1w/T2w ratios across infancy, childhood, and adolescence using a community-based neurodiverse sample. We hypothesized that volumetric and myelin-sensitive measures would exhibit distinct age-related patterns, reflecting dissociable processes of structural growth and myelin-tissue maturation. We further examined whether these developmental patterns varied as a function of biological sex. Given the inclusion of neurodevelopmentally diverse participants in our community sample, we additionally explored whether hippocampal morphology and tissue maturation differed as a function of neurodevelopmental status. In a subset of participants, psychoeducational assessments were available to characterize our sample’s cognitive functioning and behavioural adjustment.

## 2. Materials and Methods

### 2.1 Participants

Participants in this study were drawn from a community-based sample in Southwestern Ontario comprising neurotypical (NT) children and children with ASD, ADHD, or Intellectual Disability (ID). All school-age children and adolescents were attending school, and older adolescents/young adults were attending University. Participants were recruited through social media advertisements, the laboratory’s website, or the OurBrainsCAN recruitment platform. The primary inclusion criterion was being 21 years of age or younger. All participants >2 years were verbal and otherwise healthy.

Exclusion criteria included the presence of a neurological disorder or any other medical condition that could affect brain development (e.g. traumatic brain injury), as well as any MRI contraindication (i.e. pacemakers, metal in the body, etc.). For the neurodivergent group (ND), an ASD, ADHD, or ID diagnosis provided by a developmental pediatrician or an equivalent healthcare professional was needed. Additional comorbidities were permitted, given the common co-occurrence of neurodevelopmental and mental health disorders. For the NT group, participants were required to have no neurodevelopmental or mental health diagnosis and not be taking psychotropic medication. Interested parents or participants contacted the laboratory by email to express interest in participating. Participant eligibility was verified, and eligible families were provided with study information and a study brochure before a testing session was scheduled. The total sample consisted of 175 children and adolescents (74 female, 45 NT) aged between 9 months and 21 years.

The study was approved by the Health Sciences Research Ethics Board at Western University (#113515). Participants provided informed consent or assent with parental consent. All procedures were conducted in accordance with the Declaration of Helsinki.

### 2.2 Procedure

The testing session consisted of a 1-hour MRI data acquisition, conducted at the Centre for Functional and Metabolic Mapping at the Robarts Research Institute in London, Canada. A subset of children underwent a 1.5-hour cognitive testing session, which was conducted by a member of the research team in research-dedicated testing rooms at the University. Only participants aged 6 to 16 years were eligible to complete the cognitive assessment due to the age limitations of the standardized tools used (see section 2.3.1 for details).

The MRI session took place either before or after cognitive testing, depending on scanner availability, although efforts were made to conduct cognitive testing first to maximize participant performance. While participants completed the cognitive assessment, accompanying parents were asked to complete questionnaires collecting demographic information and measures of their child’s daily functioning. Completion required approximately 45 minutes on a desktop computer. If no cognitive assessment was conducted or if the parent was unavailable during the testing session, questionnaires were emailed for completion at home. For an adult participant whose parents were not involved in the research, the participant was asked to complete the demographic questionnaire, and no parental questionnaires were collected.

### 2.3 Psychoeducational measures

Participants’ cognitive abilities were assessed using the Wechsler Intelligence Scale for Children, Fifth Edition (WISC-V). Parents also completed the parent-version of the Behavior Assessment System for Children, Third Edition (BASC-3).

#### 2.3.1 WISC-V

The WISC-V is a standardized test battery that assesses the intellectual abilities of children aged 6 to 16 years (Full Scale Intelligence Quotient; FSIQ), and its five primary composite indices: the Verbal Comprehension Index (VCI), Visual Spatial Index (VSI), Fluid Reasoning Index (FRI), Working Memory Index (WMI) and Processing Speed Index (PSI). The 10 primary subtests of the battery were administered in this study, requiring approximately 75 minutes to complete. Individual assessments were conducted by a trained experimenter in a distraction-free environment. Either paper or iPad versions of the battery were administered depending on material availability, as both formats are considered equivalent (Daniel et al., 2014). Raw subtest scores were computed according to the scoring guidelines and subsequently converted into scaled scores using the English Canadian WISC-V normative sample (N=880). Scaled scores were then summed to generate composite raw scores, which were converted into standardized composite indices scores based on normative data. All WISC-V indices are standardized to a mean of 100 and a standard deviation of 15. The mean score of 100 represents average performance relative to age-matched peers, whereas higher or lower scores indicate better or poorer performance, respectively. The WISC-V is considered the gold standard for the assessment of intellectual abilities in children, and its reliability and validity are well established (Wechsler et al., 2014).

#### 2.3.2 BASC-3

The BASC-3 is a standardized questionnaire comprising a comprehensive set of scales for designed to assess the behaviors and emotions of children and adolescents. The four primary composite scales are Adaptive Skills (AS), Internalized Problems (IP), Externalized Problems (EP), and Behavioral Symptoms (BS). The parent rating forms were used in this study, including the preschool (2 to 5 years old), child (6 to 11 years old) or adolescent (12 to 21 years old) versions, depending on participant age.

Completion requires approximately 20-30 minutes, depending on the version administered. Items are rated on a 4-point Likert scale ranging from Never (0) to Almost always (3). Raw scores are summed within each scale and subsequently converted to T-scores using the U.S. normative sample (N=1800). The BASC-3 demonstrates good reliability and validity and is widely used in school settings (Reynolds & Kamphaus, 2015).

### 2.4 MRI recording and data processing

MRI scans were acquired using a 3T Siemens scanner equipped with a 32-channel head coil. A 3D-MPRAGE pulse sequence was used to acquire 192 T1-weighted slices with a thickness of 1 mm. Imaging parameters included an echo time (TE) of 2.88 ms, an inversion time of 900 ms, a flip angle of 9 degrees, a 240 × 240 matrix, and a repetition time (TR) of 2300 ms. A turbo spin-echo (TSE) sequence was used to acquire T2-weighted images with a slice thickness of 1.2 mm. Imaging parameters were as follows: TE of 109 ms, flip angle of 120 degrees, a 162 × 162 matrix, and a TR of 9000 ms.

The quality of all scans was visually assessed, and a rating from 1 (very good) to 3 (very bad) was assigned to each T1w and T2w image. T1w scans with a score of 3 (N=6) and their corresponding T2w scans were rejected from subsequent analyses.

### 2.5 Hippunfold

Hippunfold is a surface segmentation and unfolding tool for the hippocampal subfields that explicitly recognizes the cortex-like organization of the hippocampus (DeKraker et al., 2022). Briefly, it uses a deep convolutional neural network (U-Net) to segment hippocampal grey matter and surrounding tissues. Laplace fields are then applied along the anterior-posterior (AP), proximal-distal (PD), and inner-outer (IO) axes to transform each voxel into an unfolded coordinate space (DeKraker et al., 2018). Finally, a ground-truth segmentation of hippocampal subfields, derived from the 3D BigBrain histology dataset (DeKraker et al., 2020), is warped back into each participant’s native space. The resulting subfields include the subiculum (Sub), cornu ammonis (CA) 1, CA2, CA3, CA4 and dentate gyrus (DG).

Segmentation quality is assessed by visual examination and using the Dice coefficient (Dice, 1945). Within the Hippunfold pipeline, the Dice score is calculated by comparing the overlap between the whole-hippocampus mask generated by Hippunfold and the corresponding mask obtained through diffeomorphic registration to a standard template. Dice scores range from 0 to 1, with values approaching 1 indicating greater overlap.

Although segmentations generated by the two methods should be highly similar, Hippunfold is considered more anatomically precise. Accordingly, the authors recommend treating Dice scores <0.7 as failed segmentations (DeKraker et al., 2022). T1w scans with Dice scores <0.7 and visually confirmed failed segmentation (N=2) were excluded from further analyses. Subfield volumes, as well as T1w and T2w voxel intensity values, were extracted from the Hippunfold tabular outputs. Mean intensity values were computed separately for T1w and T2w images within each subfield, after which the average T1w intensity was divided by the average T2w intensity to obtain a T1w/T2w ratio. This ratio is considered a proxy measure of myelin-tissue content within brain structures (Ganzetti et al., 2014). Subfield volumes and T1w/T2w ratios were subsequently analyzed statistically.

### 2.6 Statistical analysis

All statistical analyses were performed in R (The R Foundation for Statistical Computing, version 4.6.1). First, the normality and distribution of volume and T1w/T2w ratio data were examined. Non-normal data with absolute skewness or asymmetry values >2 were winsorized at a Z score threshold of ±3.29 using the datawizard::winsorize function (v1.3.1; Patil et al., 2022). To ensure comparability across sample subgroups, demographic differences between neurodevelopment groups (ND vs NT) and sex groups were examined. Specifically, independent sample t-tests were used to assess age differences between groups, and a chi-square test was used to compare sex distributions across neurodevelopmental groups. For the subsample that completed psychoeducational testing (n=64, 37%), group differences in WISC-V FSIQ, as well as BASC-3 composite scales (AS, BS, EP and IP), were examined using t-tests. These analyses were conducted to characterize the profile of the ND and NT groups recruited from the community.

To address the study objectives, mixed-effects models were fitted using the lme4::lmer function (v2.0-1; Bates et al., 2015). In all models, age was mean-centered to facilitate interpretation and improve numerical stability. Scanner software version was included as a covariate to account for the upgrade that occurred during the study, and participant was included as a random effect to account for repeated measures.

For the primary objective, age-related patterns of hippocampal subfield volume and T1w/T2w ratio were characterized separately for each subfield (Sub, CA1, CA2, CA3, CA4, and DG). Linear, quadratic, and cubic age functions were evaluated using orthogonal polynomial terms generated with the stats::poly function (v4.6.1). Model fit was assessed using Akaike Information Criterion (AIC), Bayesian Information Criterion (BIC), and likelihood-ratio tests (stats::anova.lm v4.6.1). The optimal age-related model for each subfield-measure combination was selected based on significantly improved model fit and lower AIC and BIC values.

For the secondary objective, linear mixed-effects models were fitted using the selected age function for each subfield-measure combination. Fixed effects included age, biological sex, neurodevelopmental status, and their interactions, with hemisphere included as a control variable. Particular interest was placed on Age×Sex interactions to determine whether age-related patterns of hippocampal growth and tissue maturation differed between males and females. Neurodevelopmental status (ND vs NT) was included as an exploratory factor to evaluate whether age- and sex-related associations were robust across a neurodevelopmentally diverse sample. Type III ANOVAs were used to evaluate the significance of individual model terms. Satterthwaite’s approximation, implemented in the lmerTest package (v3.2-1; Kuznetsova et al., 2017), was used to compute test statistics. False discovery rate (FDR) correction to correct for multiple comparisons was applied across subfields using stats::p.adjust (v4.6.1). Effect sizes were estimated using partial eta-squared (effectsize::eta_squared v1.0.3; Ben-Shachar et al., 2020). Statistical significance was defined as p<0.05.

## 3. Results

### 3.1 Group demographics and psychoeducational assessments

Table 1 presents the demographic and psychoeducational characteristics of the sample. Participants in the ND group were younger than the NT group (*t*(170)=7.63, *p*=0.006).

**Table 1.** Demographic characteristics and psychoeducational assessment scores of the final sample.

| N | 172 |
| --- | --- |
| Age [mean (sd)] | 12.11 (5.05) |
| Age [range] | 9mo-21 y/o |
| Sex [N(%) F] | 74 (43.02%) |
| Dx [N(%) CTL] | 45 (26.16%) |
| BASC-3 Adaptive Skills [mean (sd)] | 39.70 (10.46) |
| BASC-3 Behavioral Symptoms [mean (sd)] | 66.33 (12.74) |
| BASC-3 Externalizing Problems [mean (sd)] | 59.98 (12.52) |
| BASC-3 Internalizing Problems [mean (sd)] | 60.29 (13.81) |
| WISC-5 Full Scale IQ [mean (sd)] | 99.75 (18.97) |
| WISC-V Verbal Comprehension [mean (sd)] | 100.02 (20.80) |
| WISC-V Visual Spatial [mean (sd)] | 101.06 (19.26) |
| WISC-V Fluid Reasoning [mean (sd)] | 103.77 (17.63) |
| WISC-V Working Memory [mean (sd)] | 98.65 (16.38) |
| WISC-V Processing Speed [mean (sd)] | 97.66 (17.02) |
| Missing or corrupted T1 scan [N(%)] | 13 (7.56%) |
| Missing or corrupted T2 scan [N(%)] | 40 (23.26%) |
| Missing BASC-3 [N(%)] | 108 (62.79%) |
| Missing WISC-V [N(%)] | 109 (63.37%) |

However, no significant differences in sex distributions were evident (*p*=0.453).

Within the subgroup for whom psychoeducational data were available (n=64, 37%), mean scores for all cognitive indices remained within the normative range for both groups (mean FSIQ: ND group=96.3; NT group=106.3). However, BASC-3 results revealed lower Adaptive Skills (*t*(59)=13.18, *p*<0.001) and higher Behavioral Symptoms (*t*(59)=11.29, *p*=0.001) in the NDD group compared to the NT group. On the WISC-V, no significant group differences in Full Scale Intelligence Quotient (*t*(59)=3.99, *p*=0.051) were evident.

No significant age differences were observed between males and females (*t*(170)=3.38, *p*=0.068), with males tending to be slightly younger than females. Within the subgroup with psychoeducational data, no significant differences in the WISC-V subscales between males and females were evident; however, only BASC-3 Adaptive Skills scores were significantly lower in females than in males (*t*(59)=5.61, *p*=0.021).

### 3.2 Age-related patterns

Tables 2 and 3 present model comparisons for hippocampal subfield volume and T1w/T2w ratio measures. Distinct age-related patterns were observed for subfield volumes and myelin-tissue processes (Fig. 1; Fig. 2). Consistent with model-selection analyses, volumes were characterized by nonlinear age-related associations, whereas T1w/T2w ratios increased linearly with age across all subfields (p_s corr_<0.05).

**Figure 1.**
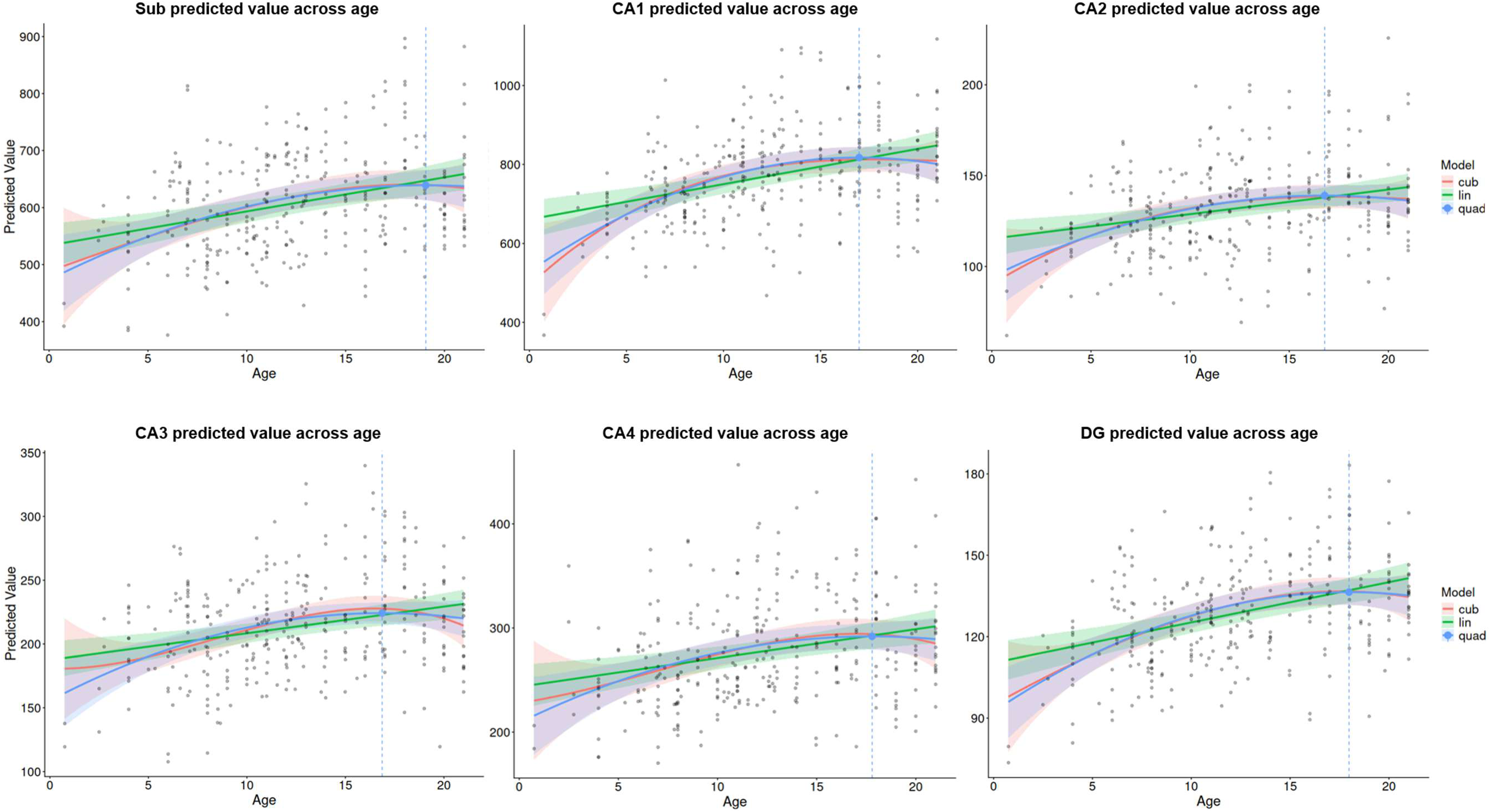
Age-related developmental patterns of hippocampal subfield volumes. *Note.* Black dots represent the observed data. Colored lines show the fitted linear, quadratic, and cubic models; shaded regions indicate their 95% confidence intervals. The blue vertical dashed line marks the estimated peak age of the quadratic model.

**Figure 2.**
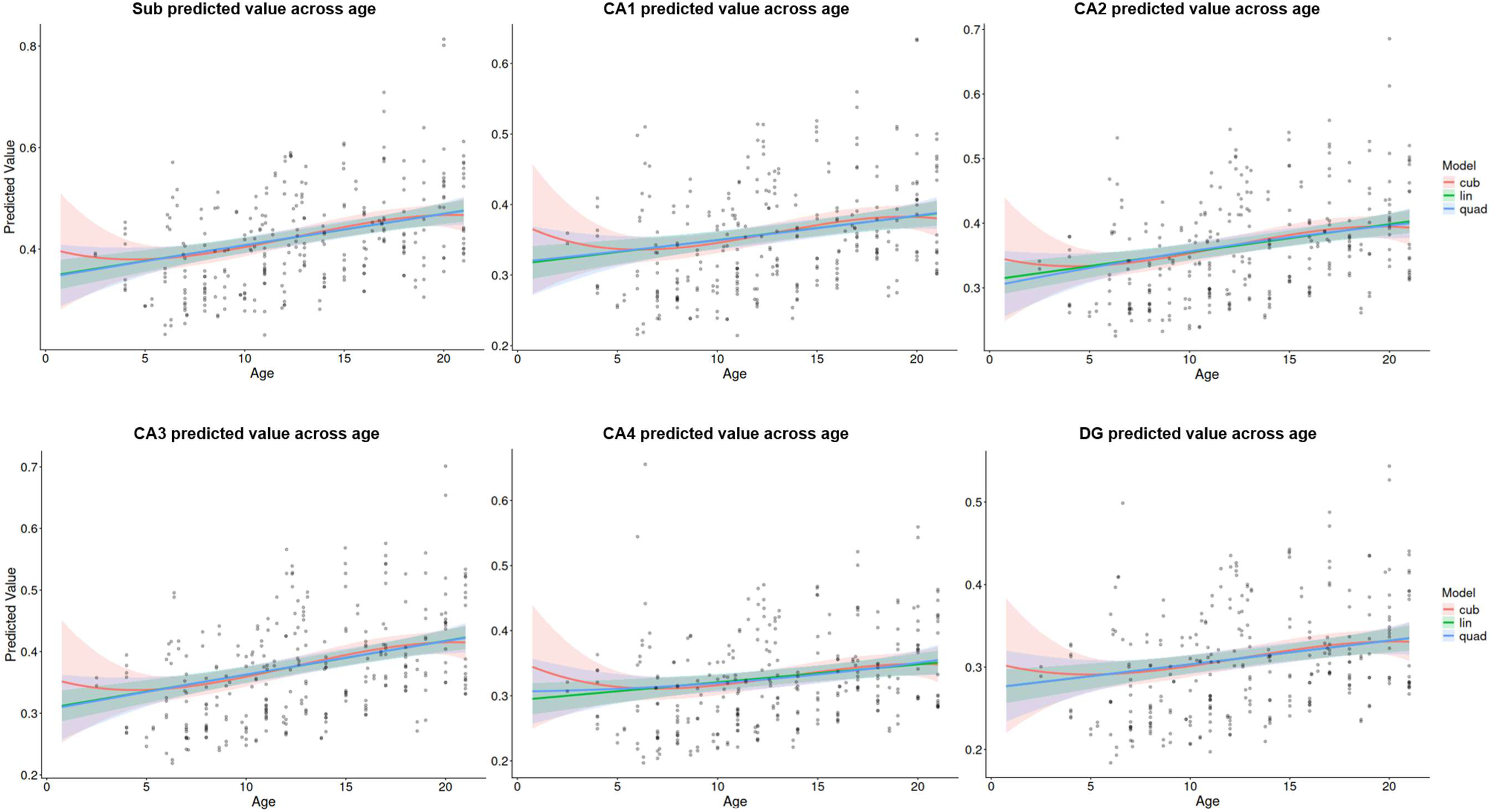
Age-related developmental patterns of hippocampal subfield T1w/T2w ratios. *Note.* Black dots represent the observed data. Colored lines show the fitted linear, quadratic, and cubic models; shaded regions indicate their 95% confidence intervals.

**Table 2.**
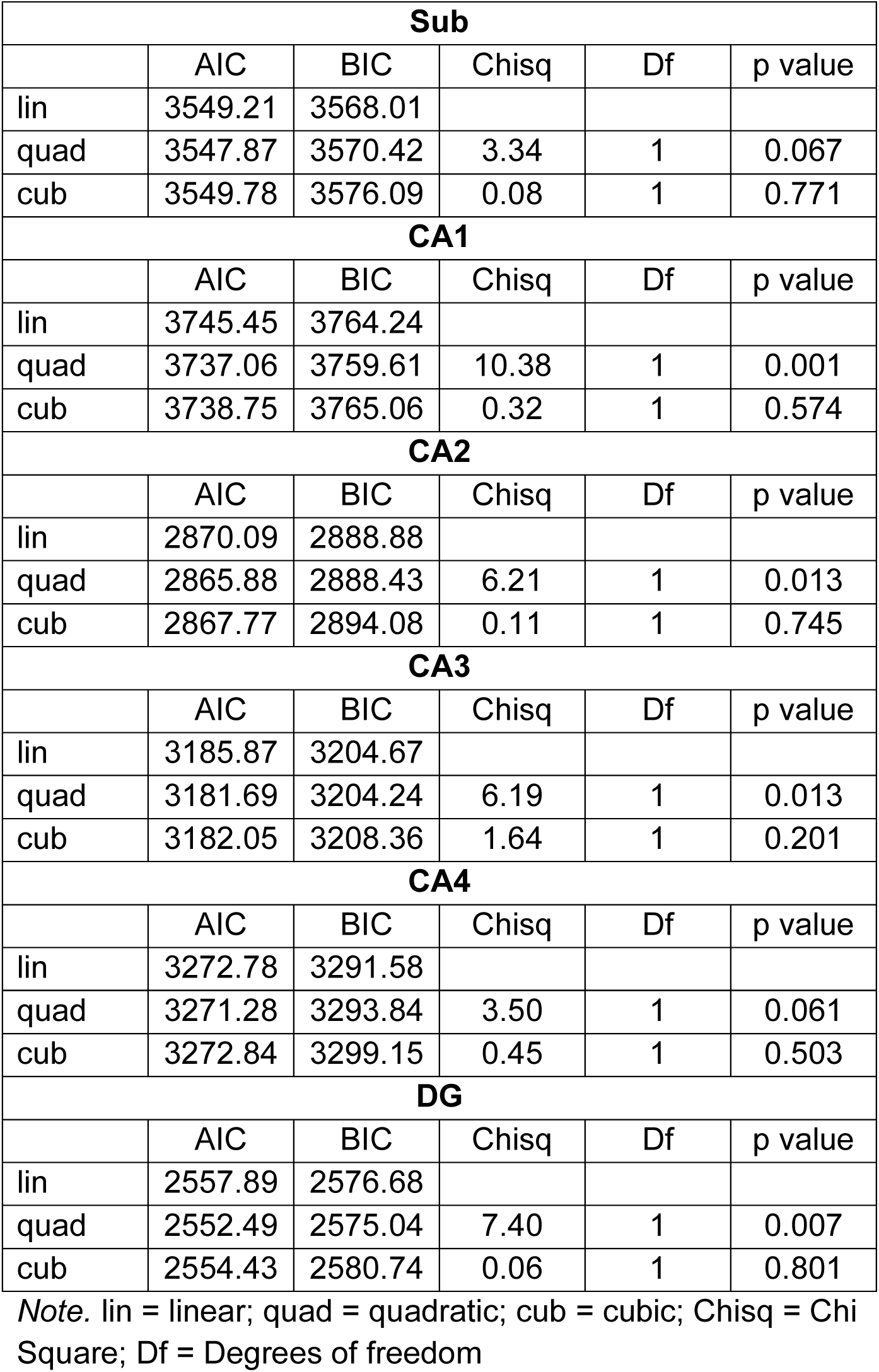
Model fit and likelihood-ratio tests of the age-related linear mixed-effect models for hippocampal subfield volumes.

| **Sub** |  |  |  |  |  |
| --- | --- | --- | --- | --- | --- |
|  | AIC | BIC | Chisq | Df | p value |
| lin | 3549.21 | 3568.01 |  |  |  |
| quad | 3547.87 | 3570.42 | 3.34 | 1 | 0.067 |
| cub | 3549.78 | 3576.09 | 0.08 | 1 | 0.771 |
| **CA1** |  |  |  |  |  |
|  | AIC | BIC | Chisq | Df | p value |
| lin | 3745.45 | 3764.24 |  |  |  |
| quad | 3737.06 | 3759.61 | 10.38 | 1 | 0.001 |
| cub | 3738.75 | 3765.06 | 0.32 | 1 | 0.574 |
| **CA2** |  |  |  |  |  |
|  | AIC | BIC | Chisq | Df | p value |
| lin | 2870.09 | 2888.88 |  |  |  |
| quad | 2865.88 | 2888.43 | 6.21 | 1 | 0.013 |
| cub | 2867.77 | 2894.08 | 0.11 | 1 | 0.745 |
| **CA3** |  |  |  |  |  |
|  | AIC | BIC | Chisq | Df | p value |
| lin | 3185.87 | 3204.67 |  |  |  |
| quad | 3181.69 | 3204.24 | 6.19 | 1 | 0.013 |
| cub | 3182.05 | 3208.36 | 1.64 | 1 | 0.201 |
| **CA4** |  |  |  |  |  |
|  | AIC | BIC | Chisq | Df | p value |
| lin | 3272.78 | 3291.58 |  |  |  |
| quad | 3271.28 | 3293.84 | 3.50 | 1 | 0.061 |
| cub | 3272.84 | 3299.15 | 0.45 | 1 | 0.503 |
| **DG** |  |  |  |  |  |
|  | AIC | BIC | Chisq | Df | p value |
| lin | 2557.89 | 2576.68 |  |  |  |
| quad | 2552.49 | 2575.04 | 7.40 | 1 | 0.007 |
| cub | 2554.43 | 2580.74 | 0.06 | 1 | 0.801 |

*Note.* lin = linear; quad = quadratic; cub = cubic; Chisq = Chi Square; Df = Degrees of freedom

**Table 3.** Model fit and likelihood-ratio tests of the age-related linear mixed-effect models for hippocampal subfield T1w/T2w ratios.

| **Sub** |  |  |  |  |  |
| --- | --- | --- | --- | --- | --- |
|  | AIC | BIC | Chisq | Df | p value |
| lin | -1075.80 | -1057.40 |  |  |  |
| quad | -1073.81 | -1051.73 | 0.01 | 1 | 0.942 |
| cub | -1072.74 | -1046.98 | 0.93 | 1 | 0.335 |
| **CA1** |  |  |  |  |  |
|  | AIC | BIC | Chisq | Df | p value |
| lin | -1140.21 | -1121.81 |  |  |  |
| quad | -1138.22 | -1116.14 | 0.01 | 1 | 0.909 |
| cub | -1137.51 | -1111.75 | 1.29 | 1 | 0.256 |
| **CA2** |  |  |  |  |  |
|  | AIC | BIC | Chisq | Df | p value |
| lin | -1107.34 | -1088.94 |  |  |  |
| quad | -1105.50 | -1083.42 | 0.16 | 1 | 0.690 |
| cub | -1104.36 | -1078.60 | 0.86 | 1 | 0.353 |
| **CA3** |  |  |  |  |  |
|  | AIC | BIC | Chisq | Df | p value |
| lin | -1154.31 | -1135.91 |  |  |  |
| quad | -1152.32 | -1130.24 | 0.01 | 1 | 0.928 |
| cub | -1151.29 | -1125.53 | 0.97 | 1 | 0.325 |
| **CA4** |  |  |  |  |  |
|  | AIC | BIC | Chisq | Df | p value |
| lin | -1078.53 | -1060.12 |  |  |  |
| quad | -1076.80 | -1054.72 | 0.27 | 1 | 0.600 |
| cub | -1075.66 | -1049.90 | 0.86 | 1 | 0.355 |
| **DG** |  |  |  |  |  |
|  | AIC | BIC | Chisq | Df | p value |
| lin | -1208.04 | -1189.64 |  |  |  |
| quad | -1206.04 | -1183.96 | 0.00 | 1 | 0.999 |
| cub | -1204.55 | -1178.79 | 0.51 | 1 | 0.475 |

*Note.* lin = linear; quad = quadratic; cub = cubic; Chisq = Chi Square; Df = Degrees of freedom

Hippocampal subfield volumes were best described by quadratic age-related associations, with lower AIC and BIC values and significant likelihood-ratio tests favoring quadratic over linear models. The only exceptions were the Sub (p=0.067) and CA4 (p=0.061), for which evidence for quadratic model fits was weaker, although the overall pattern remained consistent across subfields. In contrast, T1w/T2w ratios were best described by linear age-related associations in all subfields. More complex models did not improve model fit, and neither AIC nor BIC values supported quadratic or cubic solutions for the T1w/T2w ratio measures.

### 3.3 Final models

Tables 4 and 5 and present the final models for hippocampal subfield volume and T1w/T2w ratio measures (Fig. 3; Fig. 4). Significant main effects of age were observed for both volumes and T1w/T2w ratios (p_s corr_<0.05). Furthermore, the two measures demonstrated contrasting sex-related patterns (p_s corr_<0.05). Males exhibited larger hippocampal subfield volumes, whereas females exhibited higher T1w/T2w ratios.

**Figure 3.**
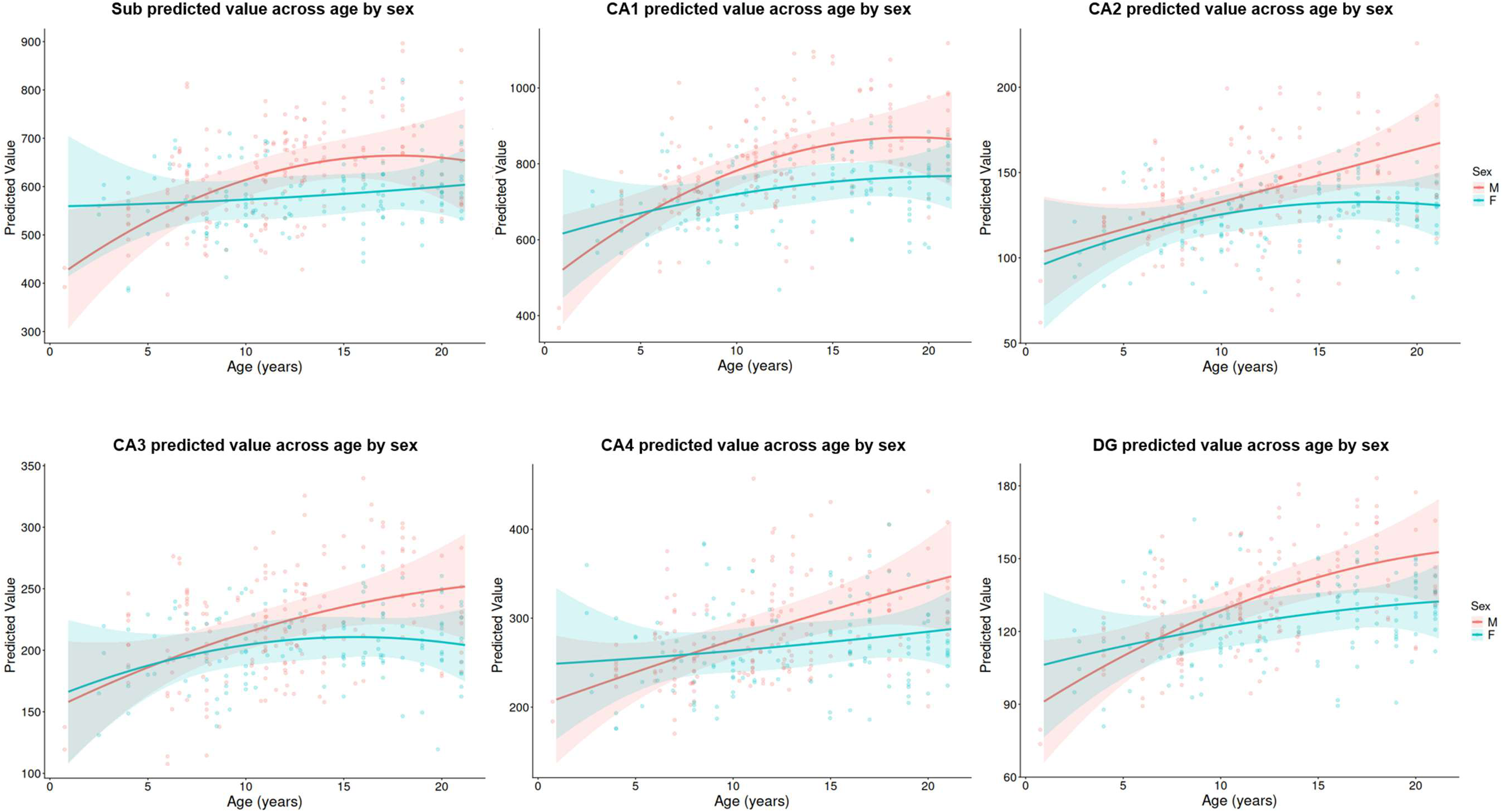
Hippocampal subfield volumes model predicted values across age by sex. *Note.* Dots represent the observed data. Colored lines show the fitted final models; shaded regions indicate their 95% confidence intervals.

**Figure 4.**
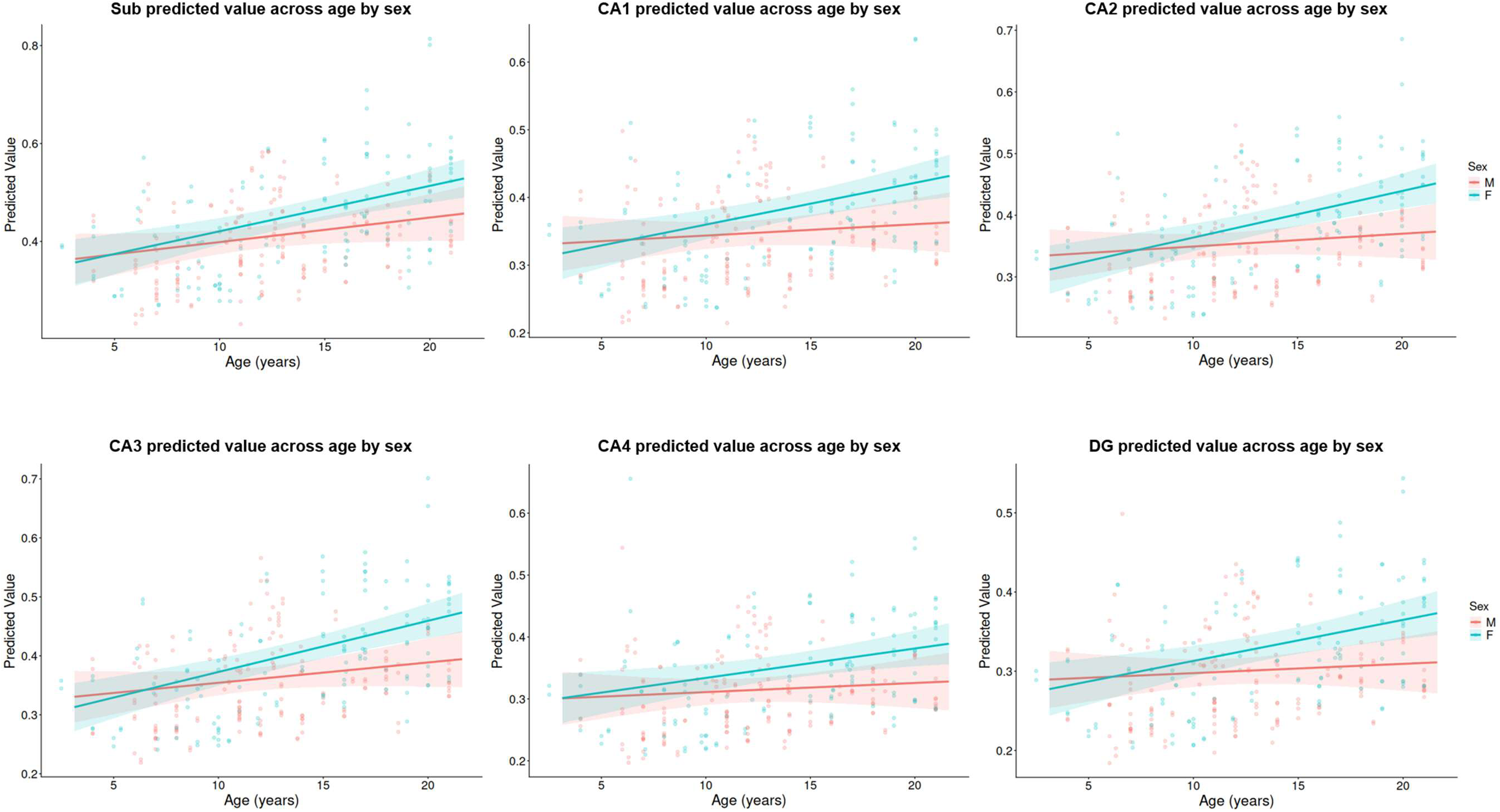
Hippocampal subfield T1w/T2w ratios model predicted values across age by sex. *Note.* Dots represent the observed data. Colored lines show the fitted final models; shaded regions indicate their 95% confidence intervals.

**Table 4.**
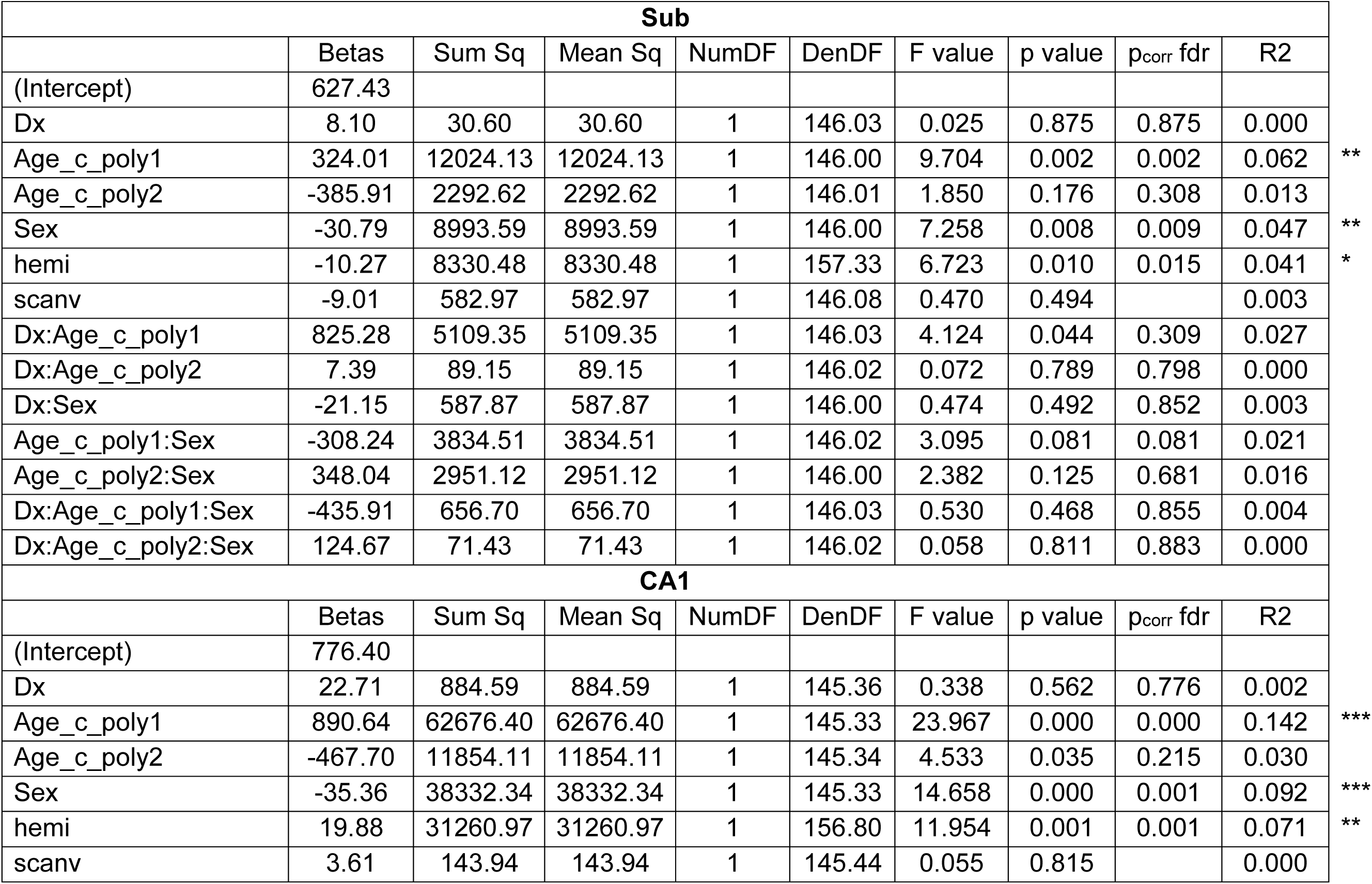
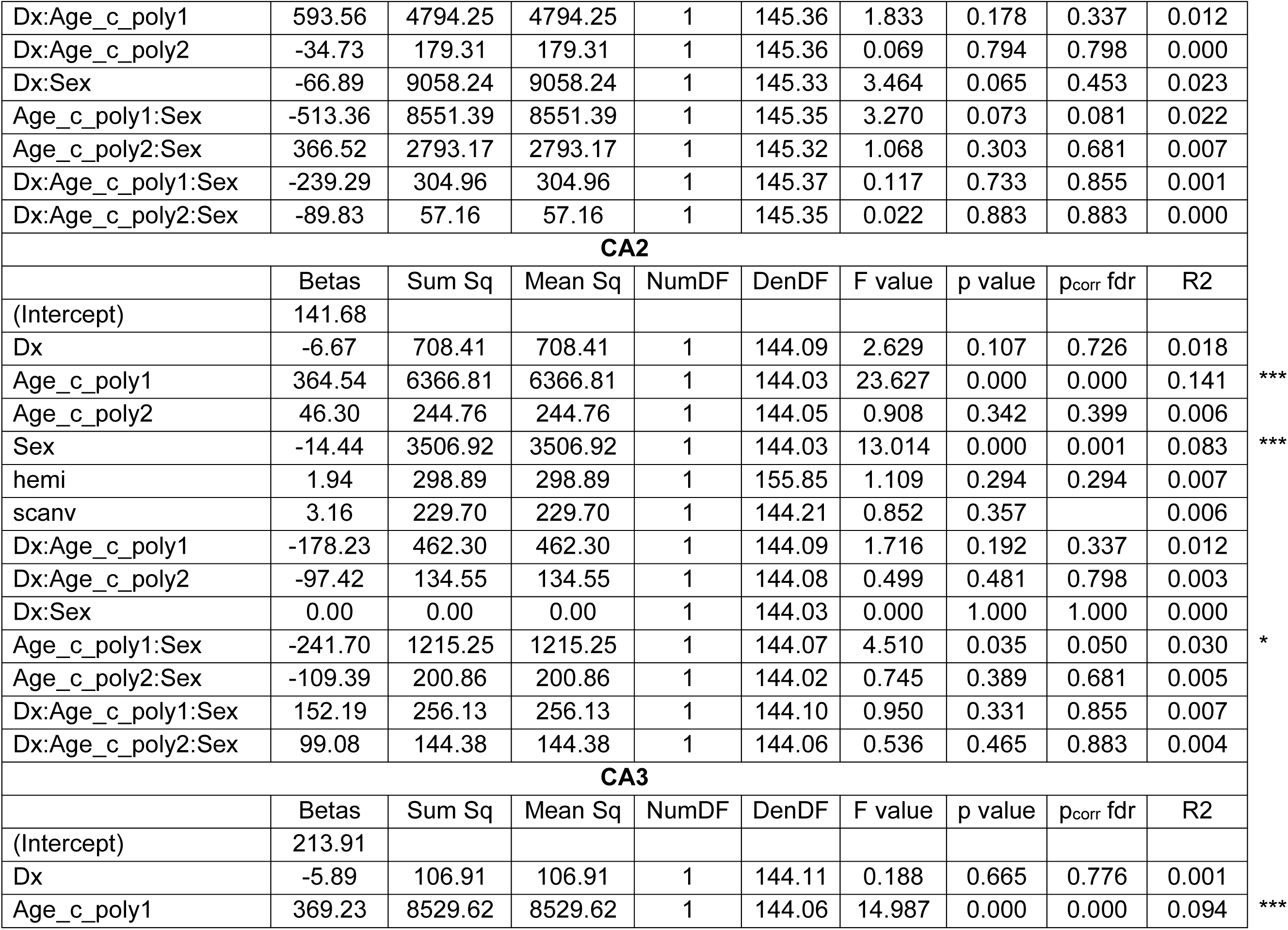
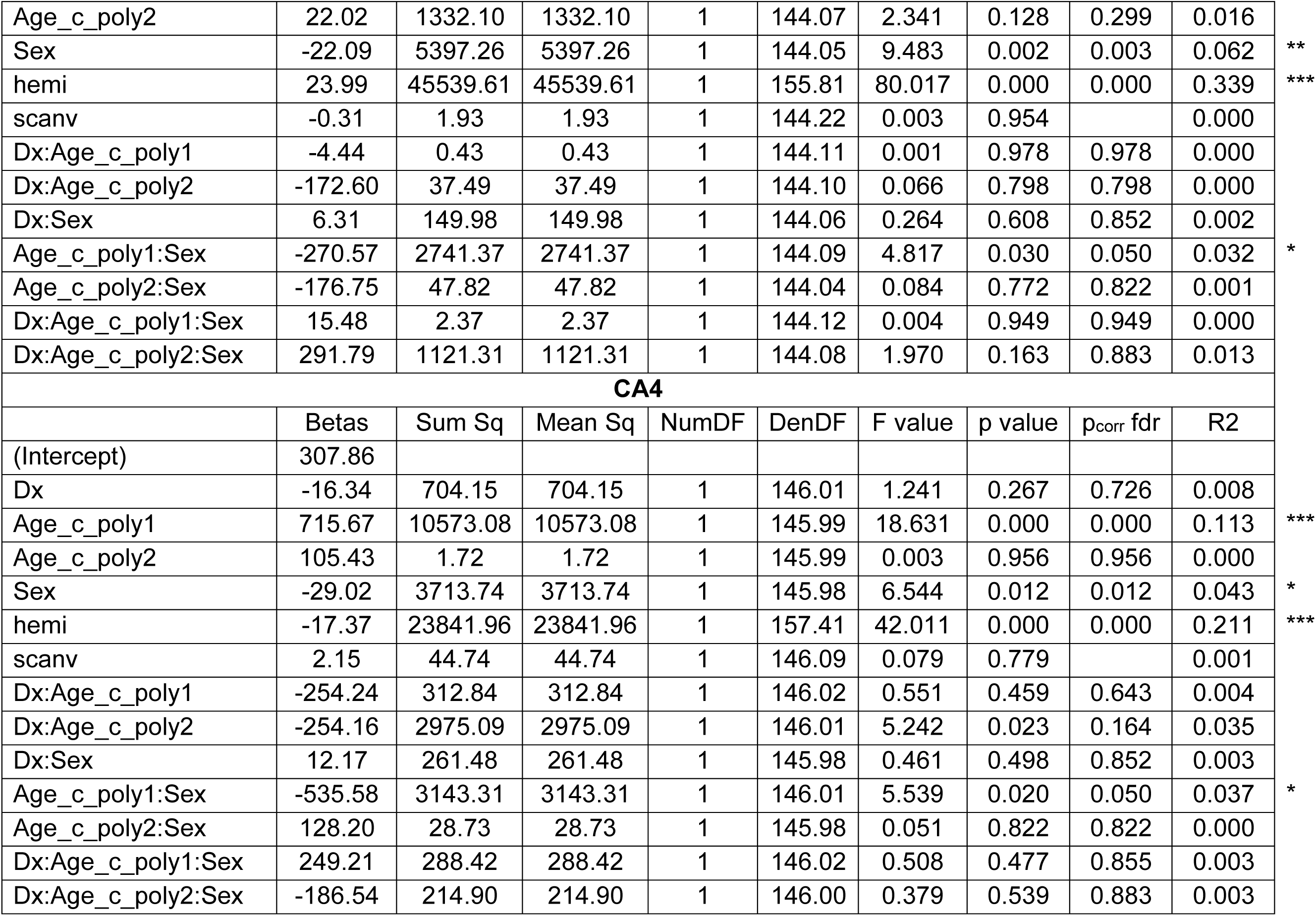
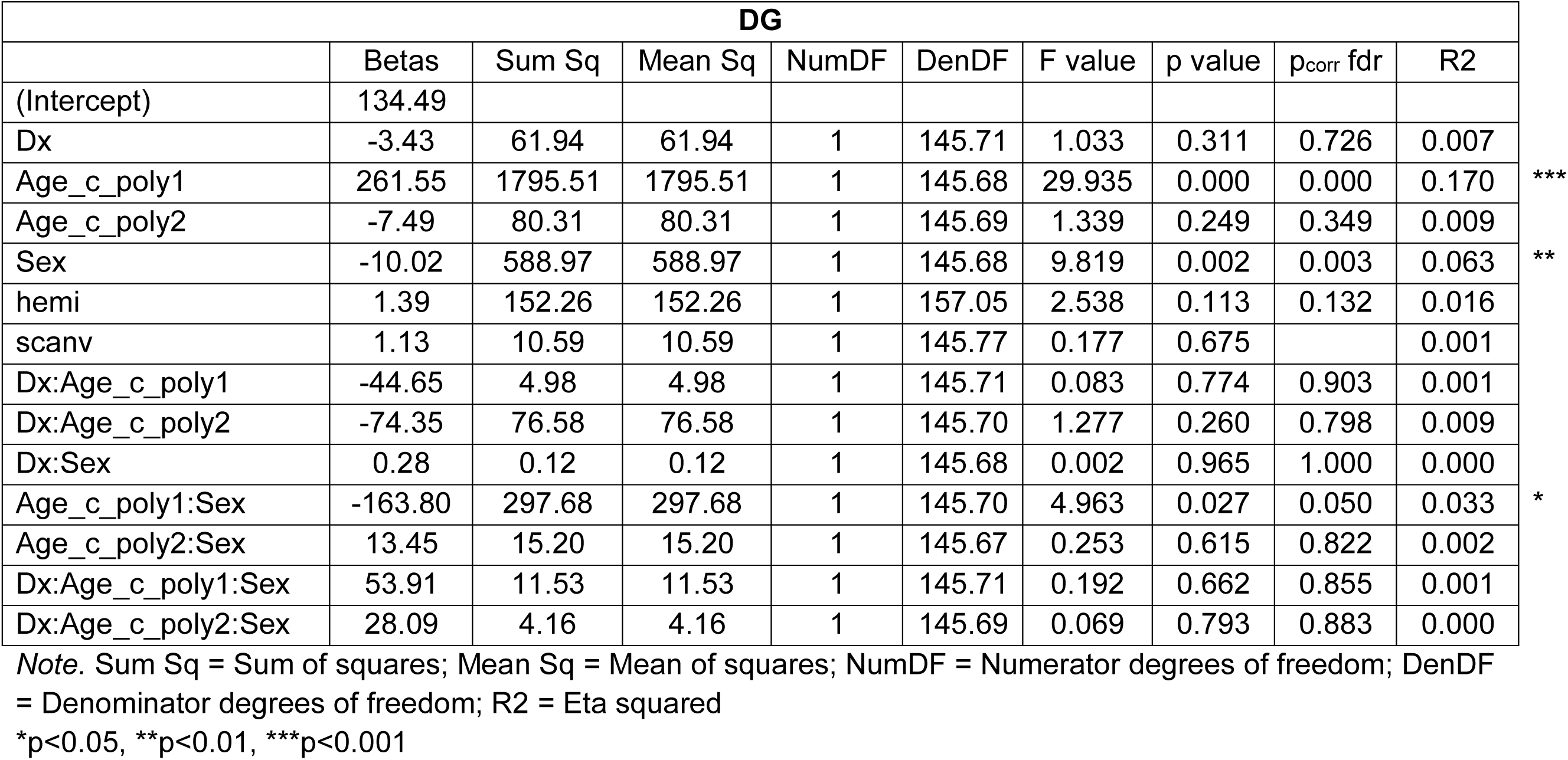
Linear mixed-effect model coefficients and ANOVA results for the hippocampal subfield volumes.

| **Sub** |  |  |  |  |  |  |  |  |  |
| --- | --- | --- | --- | --- | --- | --- | --- | --- | --- |
|  | Betas | Sum Sq | Mean Sq | NumDF | DenDF | F value | p value | p<sub>corr</sub> fdr | R2 |
| (Intercept) | 627.43 |  |  |  |  |  |  |  |  |
| Dx | 8.10 | 30.60 | 30.60 | 1 | 146.03 | 0.025 | 0.875 | 0.875 | 0.000 |
| Age_c_poly1 | 324.01 | 12024.13 | 12024.13 | 1 | 146.00 | 9.704 | 0.002 | 0.002 | 0.062 |
| Age_c_poly2 | -385.91 | 2292.62 | 2292.62 | 1 | 146.01 | 1.850 | 0.176 | 0.308 | 0.013 |
| Sex | -30.79 | 8993.59 | 8993.59 | 1 | 146.00 | 7.258 | 0.008 | 0.009 | 0.047 |
| hemi | -10.27 | 8330.48 | 8330.48 | 1 | 157.33 | 6.723 | 0.010 | 0.015 | 0.041 |
| scanv | -9.01 | 582.97 | 582.97 | 1 | 146.08 | 0.470 | 0.494 |  | 0.003 |
| Dx:Age_c_poly1 | 825.28 | 5109.35 | 5109.35 | 1 | 146.03 | 4.124 | 0.044 | 0.309 | 0.027 |
| Dx:Age_c_poly2 | 7.39 | 89.15 | 89.15 | 1 | 146.02 | 0.072 | 0.789 | 0.798 | 0.000 |
| Dx:Sex | -21.15 | 587.87 | 587.87 | 1 | 146.00 | 0.474 | 0.492 | 0.852 | 0.003 |
| Age_c_poly1:Sex | -308.24 | 3834.51 | 3834.51 | 1 | 146.02 | 3.095 | 0.081 | 0.081 | 0.021 |
| Age_c_poly2:Sex | 348.04 | 2951.12 | 2951.12 | 1 | 146.00 | 2.382 | 0.125 | 0.681 | 0.016 |
| Dx:Age_c_poly1:Sex | -435.91 | 656.70 | 656.70 | 1 | 146.03 | 0.530 | 0.468 | 0.855 | 0.004 |
| Dx:Age_c_poly2:Sex | 124.67 | 71.43 | 71.43 | 1 | 146.02 | 0.058 | 0.811 | 0.883 | 0.000 |
| **CA1** |  |  |  |  |  |  |  |  |  |
|  | Betas | Sum Sq | Mean Sq | NumDF | DenDF | F value | p value | p<sub>corr</sub> fdr | R2 |
| (Intercept) | 776.40 |  |  |  |  |  |  |  |  |
| Dx | 22.71 | 884.59 | 884.59 | 1 | 145.36 | 0.338 | 0.562 | 0.776 | 0.002 |
| Age_c_poly1 | 890.64 | 62676.40 | 62676.40 | 1 | 145.33 | 23.967 | 0.000 | 0.000 | 0.142 |
| Age_c_poly2 | -467.70 | 11854.11 | 11854.11 | 1 | 145.34 | 4.533 | 0.035 | 0.215 | 0.030 |
| Sex | -35.36 | 38332.34 | 38332.34 | 1 | 145.33 | 14.658 | 0.000 | 0.001 | 0.092 |
| hemi | 19.88 | 31260.97 | 31260.97 | 1 | 156.80 | 11.954 | 0.001 | 0.001 | 0.071 |
| scanv | 3.61 | 143.94 | 143.94 | 1 | 145.44 | 0.055 | 0.815 |  | 0.000 |
| Dx:Age_c_poly1 | 593.56 | 4794.25 | 4794.25 | 1 | 145.36 | 1.833 | 0.178 | 0.337 | 0.012 |
| Dx:Age_c_poly2 | -34.73 | 179.31 | 179.31 | 1 | 145.36 | 0.069 | 0.794 | 0.798 | 0.000 |
| Dx:Sex | -66.89 | 9058.24 | 9058.24 | 1 | 145.33 | 3.464 | 0.065 | 0.453 | 0.023 |
| Age_c_poly1:Sex | -513.36 | 8551.39 | 8551.39 | 1 | 145.35 | 3.270 | 0.073 | 0.081 | 0.022 |
| Age_c_poly2:Sex | 366.52 | 2793.17 | 2793.17 | 1 | 145.32 | 1.068 | 0.303 | 0.681 | 0.007 |
| Dx:Age_c_poly1:Sex | -239.29 | 304.96 | 304.96 | 1 | 145.37 | 0.117 | 0.733 | 0.855 | 0.001 |
| Dx:Age_c_poly2:Sex | -89.83 | 57.16 | 57.16 | 1 | 145.35 | 0.022 | 0.883 | 0.883 | 0.000 |
| **CA2** |  |  |  |  |  |  |  |  |  |
|  | Betas | Sum Sq | Mean Sq | NumDF | DenDF | F value | p value | p<sub>corr</sub> fdr | R2 |
| (Intercept) | 141.68 |  |  |  |  |  |  |  |  |
| Dx | -6.67 | 708.41 | 708.41 | 1 | 144.09 | 2.629 | 0.107 | 0.726 | 0.018 |
| Age_c_poly1 | 364.54 | 6366.81 | 6366.81 | 1 | 144.03 | 23.627 | 0.000 | 0.000 | 0.141 |
| Age_c_poly2 | 46.30 | 244.76 | 244.76 | 1 | 144.05 | 0.908 | 0.342 | 0.399 | 0.006 |
| Sex | -14.44 | 3506.92 | 3506.92 | 1 | 144.03 | 13.014 | 0.000 | 0.001 | 0.083 |
| hemi | 1.94 | 298.89 | 298.89 | 1 | 155.85 | 1.109 | 0.294 | 0.294 | 0.007 |
| scanv | 3.16 | 229.70 | 229.70 | 1 | 144.21 | 0.852 | 0.357 |  | 0.006 |
| Dx:Age_c_poly1 | -178.23 | 462.30 | 462.30 | 1 | 144.09 | 1.716 | 0.192 | 0.337 | 0.012 |
| Dx:Age_c_poly2 | -97.42 | 134.55 | 134.55 | 1 | 144.08 | 0.499 | 0.481 | 0.798 | 0.003 |
| Dx:Sex | 0.00 | 0.00 | 0.00 | 1 | 144.03 | 0.000 | 1.000 | 1.000 | 0.000 |
| Age_c_poly1:Sex | -241.70 | 1215.25 | 1215.25 | 1 | 144.07 | 4.510 | 0.035 | 0.050 | 0.030 |
| Age_c_poly2:Sex | -109.39 | 200.86 | 200.86 | 1 | 144.02 | 0.745 | 0.389 | 0.681 | 0.005 |
| Dx:Age_c_poly1:Sex | 152.19 | 256.13 | 256.13 | 1 | 144.10 | 0.950 | 0.331 | 0.855 | 0.007 |
| Dx:Age_c_poly2:Sex | 99.08 | 144.38 | 144.38 | 1 | 144.06 | 0.536 | 0.465 | 0.883 | 0.004 |
| **CA3** |  |  |  |  |  |  |  |  |  |
|  | Betas | Sum Sq | Mean Sq | NumDF | DenDF | F value | p value | p<sub>corr</sub> fdr | R2 |
| (Intercept) | 213.91 |  |  |  |  |  |  |  |  |
| Dx | -5.89 | 106.91 | 106.91 | 1 | 144.11 | 0.188 | 0.665 | 0.776 | 0.001 |
| Age_c_poly1 | 369.23 | 8529.62 | 8529.62 | 1 | 144.06 | 14.987 | 0.000 | 0.000 | 0.094 |

\*\*\*
\*\*\*
\*
\*\*\*

|  |  |  |  |  |  |  |  |  |  |  |
| --- | --- | --- | --- | --- | --- | --- | --- | --- | --- | --- |
| Age_c_poly2 | 22.02 | 1332.10 | 1332.10 | 1 | 144.07 | 2.341 | 0.128 | 0.299 | 0.016 |  |
| Sex | -22.09 | 5397.26 | 5397.26 | 1 | 144.05 | 9.483 | 0.002 | 0.003 | 0.062 | ** |
| hemi | 23.99 | 45539.61 | 45539.61 | 1 | 155.81 | 80.017 | 0.000 | 0.000 | 0.339 | *** |
| scanv | -0.31 | 1.93 | 1.93 | 1 | 144.22 | 0.003 | 0.954 |  | 0.000 |  |
| Dx:Age_c_poly1 | -4.44 | 0.43 | 0.43 | 1 | 144.11 | 0.001 | 0.978 | 0.978 | 0.000 |  |
| Dx:Age_c_poly2 | -172.60 | 37.49 | 37.49 | 1 | 144.10 | 0.066 | 0.798 | 0.798 | 0.000 |  |
| Dx:Sex | 6.31 | 149.98 | 149.98 | 1 | 144.06 | 0.264 | 0.608 | 0.852 | 0.002 |  |
| Age_c_poly1:Sex | -270.57 | 2741.37 | 2741.37 | 1 | 144.09 | 4.817 | 0.030 | 0.050 | 0.032 | * |
| Age_c_poly2:Sex | -176.75 | 47.82 | 47.82 | 1 | 144.04 | 0.084 | 0.772 | 0.822 | 0.001 |  |
| Dx:Age_c_poly1:Sex | 15.48 | 2.37 | 2.37 | 1 | 144.12 | 0.004 | 0.949 | 0.949 | 0.000 |  |
| Dx:Age_c_poly2:Sex | 291.79 | 1121.31 | 1121.31 | 1 | 144.08 | 1.970 | 0.163 | 0.883 | 0.013 |  |
| **CA4** |  |  |  |  |  |  |  |  |  |  |
|  | Betas | Sum Sq | Mean Sq | NumDF | DenDF | F value | p value | p<sub>corr</sub> fdr | R2 |  |
| (Intercept) | 307.86 |  |  |  |  |  |  |  |  |  |
| Dx | -16.34 | 704.15 | 704.15 | 1 | 146.01 | 1.241 | 0.267 | 0.726 | 0.008 |  |
| Age_c_poly1 | 715.67 | 10573.08 | 10573.08 | 1 | 145.99 | 18.631 | 0.000 | 0.000 | 0.113 | *** |
| Age_c_poly2 | 105.43 | 1.72 | 1.72 | 1 | 145.99 | 0.003 | 0.956 | 0.956 | 0.000 |  |
| Sex | -29.02 | 3713.74 | 3713.74 | 1 | 145.98 | 6.544 | 0.012 | 0.012 | 0.043 | * |
| hemi | -17.37 | 23841.96 | 23841.96 | 1 | 157.41 | 42.011 | 0.000 | 0.000 | 0.211 | *** |
| scanv | 2.15 | 44.74 | 44.74 | 1 | 146.09 | 0.079 | 0.779 |  | 0.001 |  |
| Dx:Age_c_poly1 | -254.24 | 312.84 | 312.84 | 1 | 146.02 | 0.551 | 0.459 | 0.643 | 0.004 |  |
| Dx:Age_c_poly2 | -254.16 | 2975.09 | 2975.09 | 1 | 146.01 | 5.242 | 0.023 | 0.164 | 0.035 |  |
| Dx:Sex | 12.17 | 261.48 | 261.48 | 1 | 145.98 | 0.461 | 0.498 | 0.852 | 0.003 |  |
| Age_c_poly1:Sex | -535.58 | 3143.31 | 3143.31 | 1 | 146.01 | 5.539 | 0.020 | 0.050 | 0.037 | * |
| Age_c_poly2:Sex | 128.20 | 28.73 | 28.73 | 1 | 145.98 | 0.051 | 0.822 | 0.822 | 0.000 |  |
| Dx:Age_c_poly1:Sex | 249.21 | 288.42 | 288.42 | 1 | 146.02 | 0.508 | 0.477 | 0.855 | 0.003 |  |
| Dx:Age_c_poly2:Sex | -186.54 | 214.90 | 214.90 | 1 | 146.00 | 0.379 | 0.539 | 0.883 | 0.003 |  |

| DG |  |  |  |  |  |  |  |  |  |
| --- | --- | --- | --- | --- | --- | --- | --- | --- | --- |
|  | Betas | Sum Sq | Mean Sq | NumDF | DenDF | F value | p value | p<sub>corr</sub> fdr | R2 |
| (Intercept) | 134.49 |  |  |  |  |  |  |  |  |
| Dx | -3.43 | 61.94 | 61.94 | 1 | 145.71 | 1.033 | 0.311 | 0.726 | 0.007 |
| Age_c_poly1 | 261.55 | 1795.51 | 1795.51 | 1 | 145.68 | 29.935 | 0.000 | 0.000 | 0.170 |
| Age_c_poly2 | -7.49 | 80.31 | 80.31 | 1 | 145.69 | 1.339 | 0.249 | 0.349 | 0.009 |
| Sex | -10.02 | 588.97 | 588.97 | 1 | 145.68 | 9.819 | 0.002 | 0.003 | 0.063 |
| hemi | 1.39 | 152.26 | 152.26 | 1 | 157.05 | 2.538 | 0.113 | 0.132 | 0.016 |
| scanv | 1.13 | 10.59 | 10.59 | 1 | 145.77 | 0.177 | 0.675 |  | 0.001 |
| Dx:Age_c_poly1 | -44.65 | 4.98 | 4.98 | 1 | 145.71 | 0.083 | 0.774 | 0.903 | 0.001 |
| Dx:Age_c_poly2 | -74.35 | 76.58 | 76.58 | 1 | 145.70 | 1.277 | 0.260 | 0.798 | 0.009 |
| Dx:Sex | 0.28 | 0.12 | 0.12 | 1 | 145.68 | 0.002 | 0.965 | 1.000 | 0.000 |
| Age_c_poly1:Sex | -163.80 | 297.68 | 297.68 | 1 | 145.70 | 4.963 | 0.027 | 0.050 | 0.033 |
| Age_c_poly2:Sex | 13.45 | 15.20 | 15.20 | 1 | 145.67 | 0.253 | 0.615 | 0.822 | 0.002 |
| Dx:Age_c_poly1:Sex | 53.91 | 11.53 | 11.53 | 1 | 145.71 | 0.192 | 0.662 | 0.855 | 0.001 |
| Dx:Age_c_poly2:Sex | 28.09 | 4.16 | 4.16 | 1 | 145.69 | 0.069 | 0.793 | 0.883 | 0.000 |

Note. Sum Sq = Sum of squares; Mean Sq = Mean of squares; NumDF = Numerator degrees of freedom; DenDF = Denominator degrees of freedom; R2 = Eta squared
\*p<0.05, \*\*p<0.01, \*\*\*p<0.001

**Table 5.**
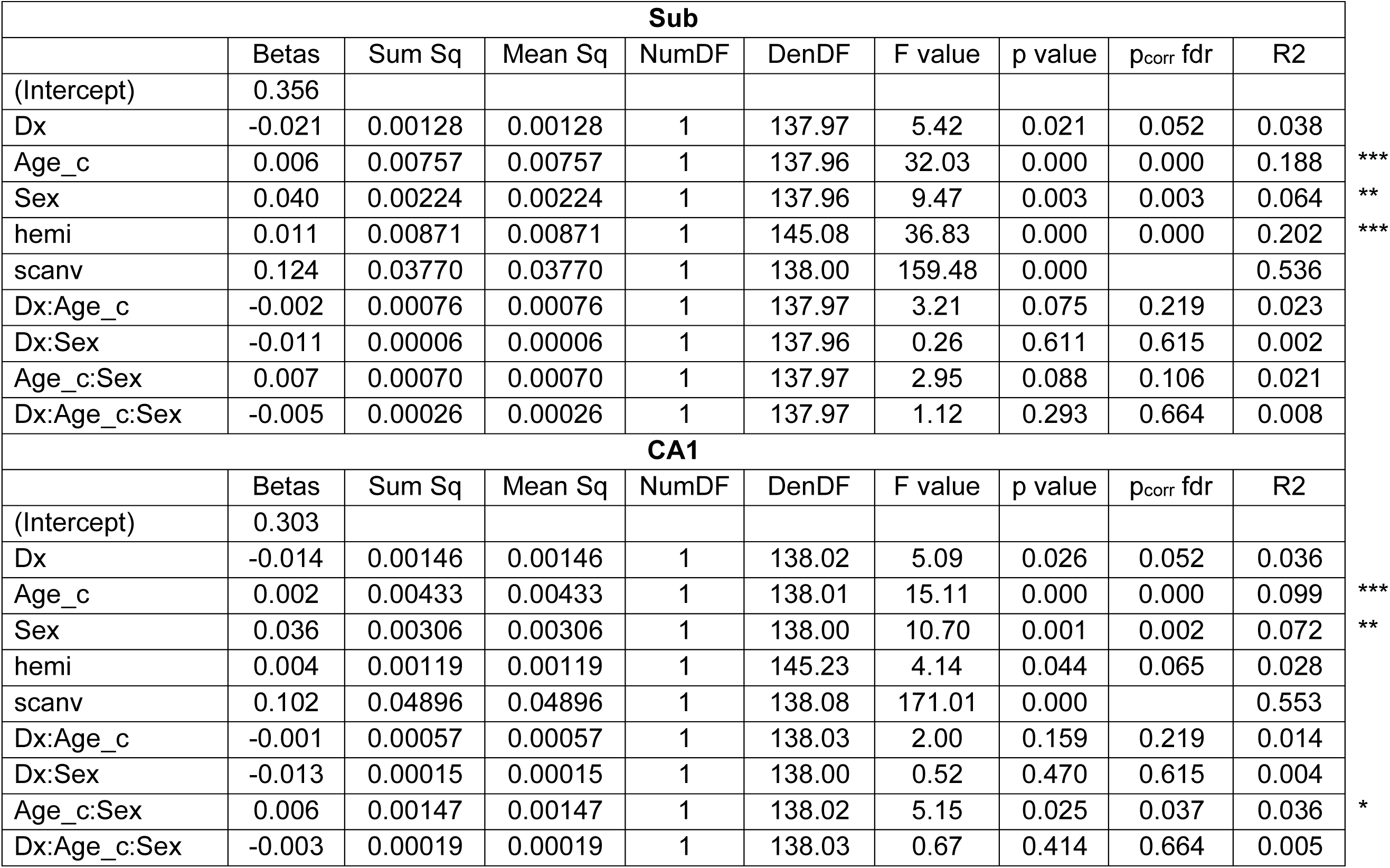
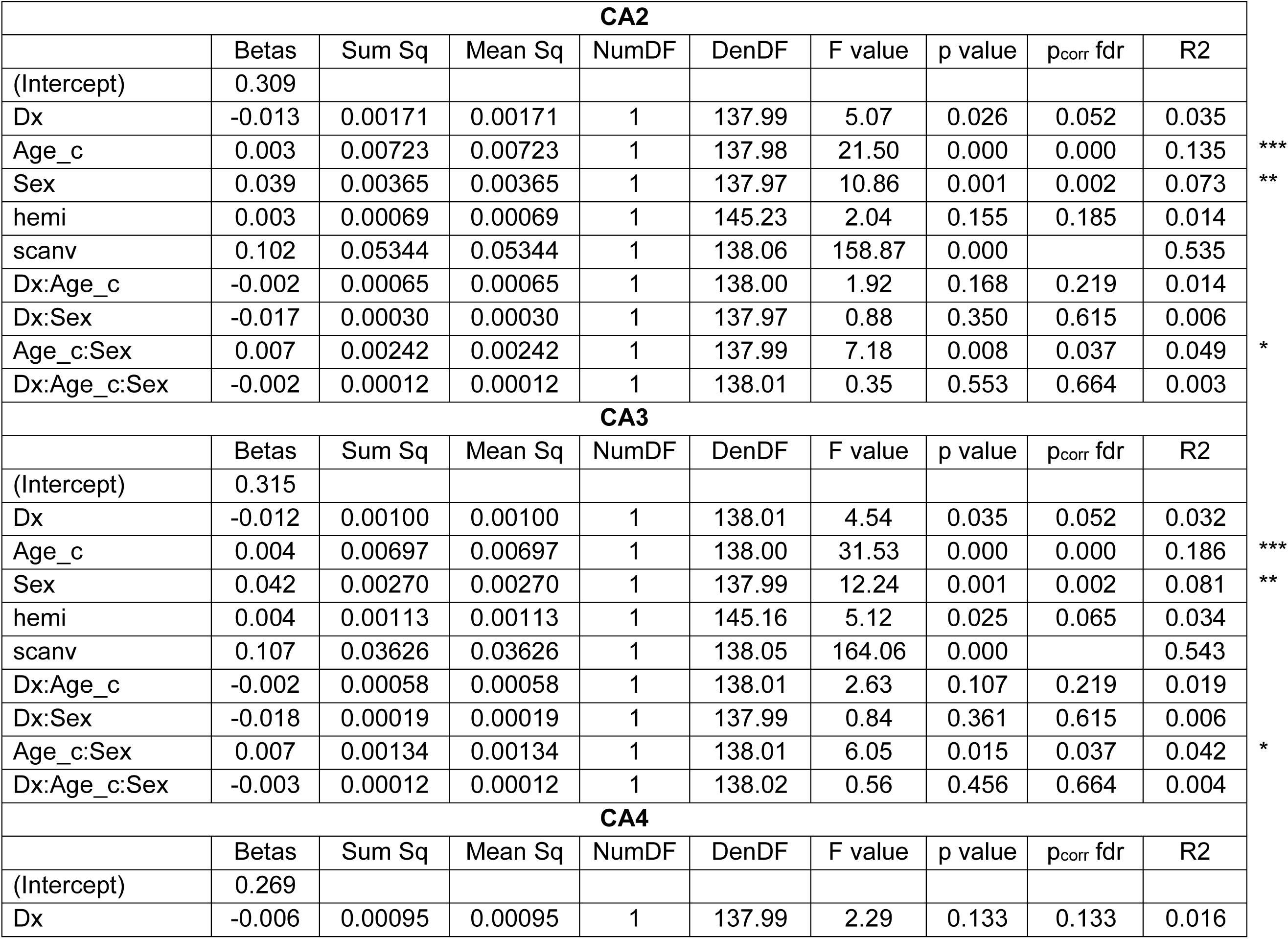
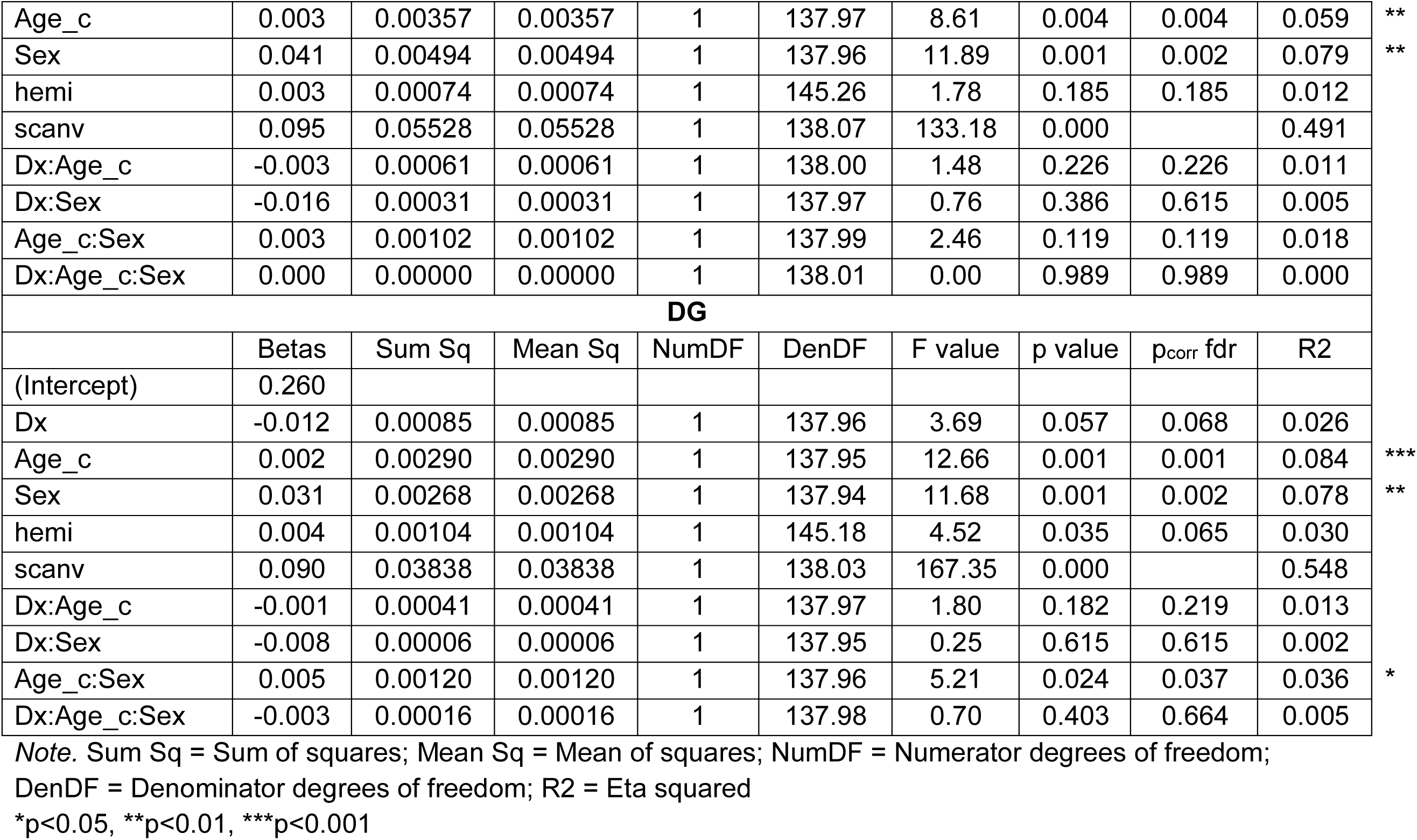
Linear mixed-effect model coefficients and ANOVA results for the hippocampal subfield T1w/Tw ratios.

| **Sub** |  |  |  |  |  |  |  |  |  |
| --- | --- | --- | --- | --- | --- | --- | --- | --- | --- |
|  | Betas | Sum Sq | Mean Sq | NumDF | DenDF | F value | p value | p<sub>corr</sub> fdr | R2 |
| (Intercept) | 0.356 |  |  |  |  |  |  |  |  |
| Dx | -0.021 | 0.00128 | 0.00128 | 1 | 137.97 | 5.42 | 0.021 | 0.052 | 0.038 |
| Age_c | 0.006 | 0.00757 | 0.00757 | 1 | 137.96 | 32.03 | 0.000 | 0.000 | 0.188 |
| Sex | 0.040 | 0.00224 | 0.00224 | 1 | 137.96 | 9.47 | 0.003 | 0.003 | 0.064 |
| hemi | 0.011 | 0.00871 | 0.00871 | 1 | 145.08 | 36.83 | 0.000 | 0.000 | 0.202 |
| scanv | 0.124 | 0.03770 | 0.03770 | 1 | 138.00 | 159.48 | 0.000 |  | 0.536 |
| Dx:Age_c | -0.002 | 0.00076 | 0.00076 | 1 | 137.97 | 3.21 | 0.075 | 0.219 | 0.023 |
| Dx:Sex | -0.011 | 0.00006 | 0.00006 | 1 | 137.96 | 0.26 | 0.611 | 0.615 | 0.002 |
| Age_c:Sex | 0.007 | 0.00070 | 0.00070 | 1 | 137.97 | 2.95 | 0.088 | 0.106 | 0.021 |
| Dx:Age_c:Sex | -0.005 | 0.00026 | 0.00026 | 1 | 137.97 | 1.12 | 0.293 | 0.664 | 0.008 |
| **CA1** |  |  |  |  |  |  |  |  |  |
|  | Betas | Sum Sq | Mean Sq | NumDF | DenDF | F value | p value | p<sub>corr</sub> fdr | R2 |
| (Intercept) | 0.303 |  |  |  |  |  |  |  |  |
| Dx | -0.014 | 0.00146 | 0.00146 | 1 | 138.02 | 5.09 | 0.026 | 0.052 | 0.036 |
| Age_c | 0.002 | 0.00433 | 0.00433 | 1 | 138.01 | 15.11 | 0.000 | 0.000 | 0.099 |
| Sex | 0.036 | 0.00306 | 0.00306 | 1 | 138.00 | 10.70 | 0.001 | 0.002 | 0.072 |
| hemi | 0.004 | 0.00119 | 0.00119 | 1 | 145.23 | 4.14 | 0.044 | 0.065 | 0.028 |
| scanv | 0.102 | 0.04896 | 0.04896 | 1 | 138.08 | 171.01 | 0.000 |  | 0.553 |
| Dx:Age_c | -0.001 | 0.00057 | 0.00057 | 1 | 138.03 | 2.00 | 0.159 | 0.219 | 0.014 |
| Dx:Sex | -0.013 | 0.00015 | 0.00015 | 1 | 138.00 | 0.52 | 0.470 | 0.615 | 0.004 |
| Age_c:Sex | 0.006 | 0.00147 | 0.00147 | 1 | 138.02 | 5.15 | 0.025 | 0.037 | 0.036 |
| Dx:Age_c:Sex | -0.003 | 0.00019 | 0.00019 | 1 | 138.03 | 0.67 | 0.414 | 0.664 | 0.005 |

\*\*\*
\*\*
\*\*\*
\*\*\*
\*\*

| CA2 |  |  |  |  |  |  |  |  |  |
| --- | --- | --- | --- | --- | --- | --- | --- | --- | --- |
|  | Betas | Sum Sq | Mean Sq | NumDF | DenDF | F value | p value | p<sub>corr</sub> fdr | R2 |
| (Intercept) | 0.309 |  |  |  |  |  |  |  |  |
| Dx | -0.013 | 0.00171 | 0.00171 | 1 | 137.99 | 5.07 | 0.026 | 0.052 | 0.035 |
| Age_c | 0.003 | 0.00723 | 0.00723 | 1 | 137.98 | 21.50 | 0.000 | 0.000 | 0.135 |
| Sex | 0.039 | 0.00365 | 0.00365 | 1 | 137.97 | 10.86 | 0.001 | 0.002 | 0.073 |
| hemi | 0.003 | 0.00069 | 0.00069 | 1 | 145.23 | 2.04 | 0.155 | 0.185 | 0.014 |
| scanv | 0.102 | 0.05344 | 0.05344 | 1 | 138.06 | 158.87 | 0.000 |  | 0.535 |
| Dx:Age_c | -0.002 | 0.00065 | 0.00065 | 1 | 138.00 | 1.92 | 0.168 | 0.219 | 0.014 |
| Dx:Sex | -0.017 | 0.00030 | 0.00030 | 1 | 137.97 | 0.88 | 0.350 | 0.615 | 0.006 |
| Age_c:Sex | 0.007 | 0.00242 | 0.00242 | 1 | 137.99 | 7.18 | 0.008 | 0.037 | 0.049 |
| Dx:Age_c:Sex | -0.002 | 0.00012 | 0.00012 | 1 | 138.01 | 0.35 | 0.553 | 0.664 | 0.003 |
| CA3 |  |  |  |  |  |  |  |  |  |
|  | Betas | Sum Sq | Mean Sq | NumDF | DenDF | F value | p value | p<sub>corr</sub> fdr | R2 |
| (Intercept) | 0.315 |  |  |  |  |  |  |  |  |
| Dx | -0.012 | 0.00100 | 0.00100 | 1 | 138.01 | 4.54 | 0.035 | 0.052 | 0.032 |
| Age_c | 0.004 | 0.00697 | 0.00697 | 1 | 138.00 | 31.53 | 0.000 | 0.000 | 0.186 |
| Sex | 0.042 | 0.00270 | 0.00270 | 1 | 137.99 | 12.24 | 0.001 | 0.002 | 0.081 |
| hemi | 0.004 | 0.00113 | 0.00113 | 1 | 145.16 | 5.12 | 0.025 | 0.065 | 0.034 |
| scanv | 0.107 | 0.03626 | 0.03626 | 1 | 138.05 | 164.06 | 0.000 |  | 0.543 |
| Dx:Age_c | -0.002 | 0.00058 | 0.00058 | 1 | 138.01 | 2.63 | 0.107 | 0.219 | 0.019 |
| Dx:Sex | -0.018 | 0.00019 | 0.00019 | 1 | 137.99 | 0.84 | 0.361 | 0.615 | 0.006 |
| Age_c:Sex | 0.007 | 0.00134 | 0.00134 | 1 | 138.01 | 6.05 | 0.015 | 0.037 | 0.042 |
| Dx:Age_c:Sex | -0.003 | 0.00012 | 0.00012 | 1 | 138.02 | 0.56 | 0.456 | 0.664 | 0.004 |
| CA4 |  |  |  |  |  |  |  |  |  |
|  | Betas | Sum Sq | Mean Sq | NumDF | DenDF | F value | p value | p<sub>corr</sub> fdr | R2 |
| (Intercept) | 0.269 |  |  |  |  |  |  |  |  |
| Dx | -0.006 | 0.00095 | 0.00095 | 1 | 137.99 | 2.29 | 0.133 | 0.133 | 0.016 |

|  |  |  |  |  |  |  |  |  |  |  |
| --- | --- | --- | --- | --- | --- | --- | --- | --- | --- | --- |
| Age_c | 0.003 | 0.00357 | 0.00357 | 1 | 137.97 | 8.61 | 0.004 | 0.004 | 0.059 | ** |
| Sex | 0.041 | 0.00494 | 0.00494 | 1 | 137.96 | 11.89 | 0.001 | 0.002 | 0.079 | ** |
| hemi | 0.003 | 0.00074 | 0.00074 | 1 | 145.26 | 1.78 | 0.185 | 0.185 | 0.012 |  |
| scanv | 0.095 | 0.05528 | 0.05528 | 1 | 138.07 | 133.18 | 0.000 |  | 0.491 |  |
| Dx:Age_c | -0.003 | 0.00061 | 0.00061 | 1 | 138.00 | 1.48 | 0.226 | 0.226 | 0.011 |  |
| Dx:Sex | -0.016 | 0.00031 | 0.00031 | 1 | 137.97 | 0.76 | 0.386 | 0.615 | 0.005 |  |
| Age_c:Sex | 0.003 | 0.00102 | 0.00102 | 1 | 137.99 | 2.46 | 0.119 | 0.119 | 0.018 |  |
| Dx:Age_c:Sex | 0.000 | 0.00000 | 0.00000 | 1 | 138.01 | 0.00 | 0.989 | 0.989 | 0.000 |  |
| **DG** |  |  |  |  |  |  |  |  |  |  |
|  | Betas | Sum Sq | Mean Sq | NumDF | DenDF | F value | p value | p<sub>corr</sub> fdr | R2 |  |
| (Intercept) | 0.260 |  |  |  |  |  |  |  |  |  |
| Dx | -0.012 | 0.00085 | 0.00085 | 1 | 137.96 | 3.69 | 0.057 | 0.068 | 0.026 |  |
| Age_c | 0.002 | 0.00290 | 0.00290 | 1 | 137.95 | 12.66 | 0.001 | 0.001 | 0.084 | *** |
| Sex | 0.031 | 0.00268 | 0.00268 | 1 | 137.94 | 11.68 | 0.001 | 0.002 | 0.078 | ** |
| hemi | 0.004 | 0.00104 | 0.00104 | 1 | 145.18 | 4.52 | 0.035 | 0.065 | 0.030 |  |
| scanv | 0.090 | 0.03838 | 0.03838 | 1 | 138.03 | 167.35 | 0.000 |  | 0.548 |  |
| Dx:Age_c | -0.001 | 0.00041 | 0.00041 | 1 | 137.97 | 1.80 | 0.182 | 0.219 | 0.013 |  |
| Dx:Sex | -0.008 | 0.00006 | 0.00006 | 1 | 137.95 | 0.25 | 0.615 | 0.615 | 0.002 |  |
| Age_c:Sex | 0.005 | 0.00120 | 0.00120 | 1 | 137.96 | 5.21 | 0.024 | 0.037 | 0.036 | * |
| Dx:Age_c:Sex | -0.003 | 0.00016 | 0.00016 | 1 | 137.98 | 0.70 | 0.403 | 0.664 | 0.005 |  |

Note. Sum Sq = Sum of squares; Mean Sq = Mean of squares; NumDF = Numerator degrees of freedom;
DenDF = Denominator degrees of freedom; R2 = Eta squared
\*p<0.05, \*\*p<0.01, \*\*\*p<0.001

Evidence for Age×Sex effects was observed for both measures, with the strongest and most consistent interactions occurring for T1w/T2w ratios. Age×Sex interactions for T1w/T2w ratios were observed in CA1 (p_corr_=0.037), CA2 (p_corr_=0.037), CA3 (p_corr_=0.037), and DG (p_corr_=0.037), indicating steeper age-related increases in tissue maturation in females than in males. For volume measures, corresponding effects trended towards significance and localized to CA2 (p_corr_=0.050), CA3 (p_corr_=0.050), CA4 (p_corr_=0.050), and DG (p_corr_=0.050), with non-significant effects observed in the Sub and CA1 (p_corr_=0.081).

Significant main effects of hemisphere were observed in several hippocampal volumetric subfields. For volume measures, the Sub (p_corr_=0.015) and CA4 exhibited larger volumes in the left hemisphere than in the right, whereas CA1 (p_corr_=0.001) and CA3 (p_corr_< 0.001) exhibited larger volumes in the right hemisphere than in the left. For T1w/T2w ratios, the Sub showed significantly higher values in the right hemisphere (p_corr_<0.001). Similar rightward asymmetries were observed for CA1 (p_corr_=0.067), CA3 (p_corr_=0.065), and DG (p_corr_=0.065), although these effects did not survive correction for multiple comparisons.

Neurodevelopmental status was not significantly associated with hippocampal subfield volume or T1w/T2w ratio following correction for multiple comparisons. Trends of lower T1w/T2w ratios was observed in the ND participants across the Sub (p_corr_=0.052), CA1 (p_corr_=0.052), CA2 (p_corr_=0.052), CA3 (p_corr_=0.052), and DG (p_corr_=0.068) compared to the NT group.

## 4. Discussion

We examined age-related variation in hippocampal subfield volumes and T1w/T2w ratios in a community-based sample of infants, children, and adolescents spanning 9 months to 21 years of age. Hippocampal subfields’ macrostructure and microstructure exhibited distinct age-related patterns. Whereas hippocampal macrostructural volumes were best characterized by nonlinear associations with age, T1w/T2w ratios increased linearly across the developmental age range. More specifically, hippocampal volumes increased throughout childhood before reaching a peak and subsequently stabilizing, whereas T1w/T2w ratios continued to increase across the age range studied. These findings suggest that structural growth and tissue maturation represent partially dissociable biological processes during hippocampal development.

### 4.1 Age-related patterns of hippocampal growth and maturation

Our findings demonstrating that hippocampal subfield growth is quadratic, but subfield maturation is linear are supported by the broader developmental imaging literature. Linear increases in white matter and cortical myelination have been reported as well as inverted U-shaped patterns in grey matter with increases in childhood and further decreases in adolescence (Grydeland et al., 2013; Houston et al., 2014; Mills et al., 2016; Norbom et al., 2020; Shafee et al., 2015; Sussman et al., 2016). Our findings support previous reports of non-linear age-related changes in hippocampal subfield volumes throughout childhood and adolescence (Krogsrud et al., 2014; Lee et al., 2014; Mu et al., 2020; Sussman et al., 2016), but contrast with studies reporting linear, negative or null associations (Karat et al., 2024; Tamnes et al., 2018). Additional studies with larger samples and longitudinal designs are needed to establish patterns of hippocampal subfield development more conclusively.

The differing age-related patterns observed for volume and T1w/T2w measures may reflect distinct underlying neurobiological mechanisms. Volumetric development likely reflects processes associated with structural growth, including cellular proliferation, dendritic arborization, and the establishment of hippocampal architecture. In contrast, T1w/T2w ratios are thought to reflect tissue composition and myelination (Ganzetti et al., 2014) and may therefore capture ongoing processes in development that continue throughout childhood and adolescence. Consistent with this interpretation, age-by-sex interaction effects were most pronounced for T1w/T2w measures, with females exhibiting steeper age-related increases in tissue maturation across multiple hippocampal subfields despite largely comparable cognitive and behavioural profiles. Findings indicate that hippocampal development cannot be fully characterized by morphology alone and that microstructural measures reflective of myelination may provide complementary information regarding developmental timing and biological maturity.

In contrast to several previous studies reporting differential neurodevelopmental patterns across hippocampal subfields (Lee et al., 2014, 2020; Mu et al., 2020; Sussman et al., 2016; Tamnes et al., 2018), we observed remarkably similar age-related patterns across subfields. However, our findings are consistent with those of Krogsrud and colleagues (2014) and a meta-analysis addressing this question, which likewise reported relatively homogeneous developmental associations across subfields (Homayouni et al., 2023). Differences in segmentation protocols, model-selection approaches, and statistical complexity may contribute substantially to discrepancies across studies. Our use of Hippunfold and parsimonious model-selection procedures may have enhanced sensitivity to common developmental patterns across subfields.

Interestingly, neurodevelopmental status was not significantly associated with hippocampal subfield volume or T1w/T2w ratios following correction for multiple comparisons. Although trend-level reductions in T1w/T2w ratios were observed across several subfields in ND participants, the absence of neurodevelopmental effects suggests that the age- and sex-related patterns observed here are largely preserved across a broad range of developmental presentations. This finding may reflect the community-based nature of the sample, which included neurodevelopmentally diverse participants who were all verbal and high functioning. Future studies with larger sample sizes will be required to determine whether specific neurodevelopmental conditions are associated with alterations in hippocampal tissue maturation that are not apparent at the morphological level.

We also observed evidence of hemispheric asymmetry within hippocampal subfields. Consistent with previous reports, CA1 and CA3 exhibited rightward volumetric asymmetry (Krogsrud et al., 2014; Mu et al., 2020; Schmidt et al., 2018; Uematsu et al., 2012), whereas the Sub and CA4 were larger in the left hemisphere. T1w/T2w ratios also showed a predominantly rightward pattern, with higher values observed across all right hippocampal subfields and significant effects in the Sub. Findings support emerging evidence that hippocampal lateralization varies across subfields. Given the distinct functional roles of the left and right hippocampus (Nichols et al., 2023; Persson & Söderlund, 2015), as well as their differential involvement in neurodevelopmental and psychiatric disorders (Deniz et al., 2026; Haukvik et al., 2018; Sun et al., 2023), further research is needed to clarify the biological significance of hippocampal asymmetric developmental patterns.

### 4.2 Sex-specific patterns of hippocampal growth and maturation

We identified robust sex differences in hippocampal subfields. The absence of corresponding sex differences in intellectual functioning and the limited behavioural differences observed in the sample suggest that these findings are unlikely to be explained by cognitive performance alone. Instead, they may reflect differences in the timing or rate of hippocampal development.

Males exhibited larger hippocampal subfield volumes, consistent with previous studies reporting larger whole-hippocampus volumes in males than in females from as early as the third trimester of gestation (Fish et al., 2020; Nichols et al., 2024; Ruigrok et al., 2014; Tan et al., 2016; Wierenga et al., 2014). This makes sense given the larger overall brain volumes typically observed in males (Kaczkurkin et al., 2019; Ruigrok et al., 2014). Studies examining the hippocampal subfields volumes have likewise generally reported larger volumes in males across most subfields (Krogsrud et al., 2014; Mu et al., 2020; Riggins et al., 2018; Tamnes et al., 2018), although contrary findings have also been reported (Sussman et al., 2016). As noted by Tan and colleagues (2016), some of this variability may be attributable to differences in whether hippocampal volumes are adjusted for total brain volume, as such corrections can attenuate or eliminate apparent sex differences. Nevertheless, meta-analytic evidence consistently supports larger uncorrected hippocampal volumes in males (Ruigrok et al., 2014; Tan et al., 2016), making our findings broadly consistent with the existing literature.

We also observed significant sex differences in developmental trajectories, with males showing steeper age-related increases in hippocampal volume than females. Such interactions have rarely been examined in studies of hippocampal subfields. Findings similar to ours (Karat et al., 2024; Mu et al., 2020), as well as conflicting with ours (Riggins et al., 2018; Sussman et al., 2016; Tamnes et al., 2018) have been reported. Studies examining the whole hippocampus during childhood and adolescence have also reported steeper age-related volumetric increases in males (Fish et al., 2020; Herting et al., 2018; Nichols et al., 2024), but inconsistent findings are common in lifespan studies (Narvacan et al., 2017; Tan et al., 2016; Wierenga et al., 2014), suggesting that age-by-sex interactions may become less pronounced in adulthood and later life. These sex differences in hippocampal development may contribute to our understanding of sex differences in the prevalence of neurodevelopmental and psychiatric disorders.

Normative sex differences in hippocampal morphology (males > females) may represent an important factor contributing to differential vulnerability across disorders. For example, depression, bipolar disorder, and Alzheimer’s disease disproportionately affect females and have been associated with reduced hippocampal volumes (Hibar et al., 2016; Rao et al., 2023; Schmaal et al., 2016). In contrast, ASD is more prevalent in males and has been associated with larger hippocampal volumes (Deniz et al., 2026).

In contrast to volumetric findings, females exhibited higher T1w/T2w ratios and steeper age-related increases in this measure than males. This question has received limited attention to date, but two recent studies reported higher T1w/T2w ratios across cortical and subcortical regions in males than in females (Corrigan et al., 2021; Norbom et al., 2020). Although the hippocampus was not examined specifically in those studies, they nevertheless suggest evidence of sexual dimorphism in brain myelination. One particularly relevant study, using magnetization transfer (MT) a measure even more specific to myelin content than a T1w/T2w ratio, reported higher hippocampal myelin-sensitive MT values in females than in males in supplementary analyses (Ziegler et al., 2019). Given myelination’s role in constraining neuroplasticity and facilitating efficient information transfer (Glasser et al., 2014), these findings could indicate greater resilience to the effects of adverse experiences and environmental influences in females.

### 4.3 Limitations

While this study has many strengths, a few limitations are noted here. First, the cross-sectional design means that the developmental patterns reported here are inferred from comparisons across individuals of different ages rather than from within-individual changes over time. We employed a novel segmentation tool to examine the macrostructure of hippocampal subfields using a specific proximal-distal/transverse axis segmentation protocol. However, longitudinal axis segmentation protocols also exist and may provide complementary information, given their demonstrated functional relevance (Poppenk et al., 2013; Strange et al., 2014). Further studies could also integrate both approaches, as some authors have suggested (Fanselow & Dong, 2010; Genon et al., 2021), for example by using a surface vertex based approach to obtain a more comprehensive and fine-grained characterization of hippocampal structure and function. Such surface-based analyses may also enable the investigation of additional morphometric features, including hippocampal thickness and gyrification, which could provide greater sensitivity to localized developmental changes than regional volume measures alone. Finally, this study focused on structural variations in hippocampal subfields. Additional research will be required to determine the functional implications of these developmental and neurodivergence-related differences.

## 5. Conclusion

Here, we examined age-related variation in hippocampal subfield volumes and tissue maturation reflective of myelination processes in a community-based sample across a broad age range. Hippocampal subfield structural growth and tissue maturation followed distinct age-related patterns, with nonlinear changes in volume but prolonged linear increases in tissue myelin revealed through T1w/T2w ratios, which were not attributed to differences in psychoeducational profiles. Further, females exhibited steeper age-related increases in tissue maturation compared to males despite largely comparable cognitive and behavioural profiles. Findings suggest that structural growth and tissue maturation represent partially dissociable biological processes during human hippocampal development and that measures of tissue composition may capture developmental variation not apparent from morphology alone.

## 6. Declaration of the use of AI

AI (or other tools) were used to reword and check spelling and grammar of our draft text.

## 7. Data and code availability

The datasets analysed and the code generated during the current study are available from the corresponding author on reasonable request.

## 8. Author contributions

K.R. took part in conceptualization and developing the methodology of the study, performing the data collection and curation, did the analyses and wrote the initial draft of the manuscript.

E.S.N. took part in conceptualization and developing the methodology of the study and reviewed and edited the manuscript.

M.S. and A.R.K. did the software development and provided support on its usage, as well as reviewed and edited the manuscript

E.G.D. took part in conceptualization and the development of the methodology of the study, acquired the funding and provided the resources to carry out the project, reviewed and edited the manuscript and provided supervision throughout the execution of the project.

